# Exploring how ecological and epidemiological processes shape multi-host disease dynamics using global sensitivity analysis

**DOI:** 10.1101/2022.11.01.514725

**Authors:** Kalpana Hanthanan Arachchilage, Mohammed Y. Hussaini, N. G. Cogan, Michael H. Cortez

**Affiliations:** Department of Mathematics, Florida State University, Tallahassee Fl 32306; Department of Biological Science, Florida State University, Tallahassee Fl 32306

**Keywords:** global sensitivity analysis, dilution effect, Partial Rank Correlation Coefficient, environmental transmission, Daphnia

## Abstract

We use global sensitivity analysis (specifically, Partial Rank Correlation Coefficients) to explore the roles of ecological and epidemiological processes in shaping the temporal dynamics of a parameterized SIR-type model of two host species and an environmentally transmitted pathogen. We compute the sensitivities of disease prevalence in each host species to model parameters. Sensitivity rankings and subsequent biological interpretations are calculated and contrasted for cases were the pathogen is introduced into a disease-free community and where a second host species is introduced into an endemic single-host community. In some cases the magnitudes and dynamics of the sensitivities can be predicted only by knowing the host species characteristics (i.e., their competitive abilities and disease competence) whereas in other cases they can be predicted by factors independent of the species characteristics (specifically, intraspecific versus interspecific processes or the species’ roles of invader versus resident). For example, when a pathogen is initially introduced into a disease-free community, disease prevalence in both hosts is more sensitive to the burst size of the first host than the second host. In comparison, disease prevalence in each host is more sensitive to its own infection rate than the infection rate of the other host species. In total, this study illustrates that global sensitivity analysis can provide useful insight into how ecological and epidemiological processes shape disease dynamics and how those effects vary across time and system conditions. Our results show that sensitivity analysis can provide quantification and direction when exploring biological hypotheses.

## 1 Introduction

Many empirical studies have shown that the absence or presence of a second host species can increase or decrease levels of disease in a focal host species (Telfer et al., 2005; Dizney and Ruedas, 2009; Searle et al., 2016; Levine et al., 2017; Hydeman et al., 2017; Zimmermann et al., 2017; Luis et al., 2018). This empirical work is complemented by modeling studies exploring how specific processes shape disease dynamics in two-host communities (Rudolf and Antonovics, 2005; O’Regan et al., 2015; Searle et al., 2016; Roberts and Heesterbeek, 2018; Cortez and Duffy, 2021). Combined, these studies indicate that changes in focal host disease levels depend on both ecological processes (such as intraspecific and interspecific competition between host species for resources) and epidemiological processes (such as transmission, recovery, and disease-induced mortality). For example, disease levels in a focal host often increase when the second host has high competence (i.e., a high ability to transmit the disease), however disease levels in the focal host can instead decrease if interspecific competition between the host species is sufficiently strong (O’Regan et al., 2015; Searle et al., 2016; Cortez and Duffy, 2021). In combination with empirical and theoretical work on communities with more than two host species (Dobson, 2004; Roche et al., 2012; Joseph et al., 2013; Mihaljevic et al., 2014; Faust et al., 2017; Cortez, 2021), the effects of host species richness of disease dynamics are likely to be context-dependent and depend on the specific characteristics of the species present in the community (LoGiudice et al., 2008; Randolph and Dobson, 2012; Rohr et al., 2020; Halliday et al., 2020). This study uses global sensitivity analysis to investigate the roles of ecological and epidemiological processes in shaping the temporal dynamics of a two-host epidemiological model.

The existing body of theory on multi-host communities has been fruitful in identifying some of the ways in which ecological and epidemiological processes shape disease-dynamics in multi-host communities. However, a key limitation of nearly every study is that they focus on asymptotic regimes of the models. In particular, many studies (e.g., Dobson 2004; O’Regan et al. 2015; Roberts and Heesterbeek 2018) focus on the pathogen basic reproduction number 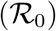, which is a measure of how fast the density or proportion of infected hosts in a population will increase in the limit where the pathogen is rare. Many other studies (e.g., Rudolf and Antonovics 2005; Roberts and Heesterbeek 2018; Cortez and Duffy 2021; Cortez 2021) focus on the density or the proportion of infected individuals in a focal host at equilibrium; these studies focus on disease dynamics in the limit where time is unbounded (*t* → ∞). Importantly, these metrics can disagree (Roche et al., 2012; Roberts and Heesterbeek, 2018; Cortez and Duffy, 2021). For example, increased transmission by a second host can increase 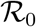 while also decreasing the proportion of infected individuals at equilibrium (Cortez and Duffy, 2021). The disagreement between metrics implies that the effects of at least one ecological or epidemiological process must differ in sign or magnitude between the two asymptotic regimes. Unfortunately, because current theory only focuses on asymptotic regimes of multi-host models, it is limited in its ability to predict when and why the effects of specific ecological or evolutionary processes change over time. Thus, new theory is needed to help explain the different ways in which ecological and epidemiological processes shape disease dynamics over different time scales.

From a mathematical perspective, understanding the mechanisms or characteristics that have the most influence on specific outcomes (such as the proportion of infected individuals or other measures of disease dynamics) requires insight into the roles of specific parameters within a predictive model – for which sensitivity analysis is precisely designed. In this regard we can interpret a deterministic model with uncertain parameter inputs as a form of stochasticity and quantify how uncertainty in the parameters propagates through the model and affects model dynamics.

The ingredients for sensitivity analysis are:

- Input parameters (*γ_i_*) – these may be considered as stochastic parameters with a given distribution.
- A model – some relationship between input parameters and an output. Generally we consider a system of differential equations

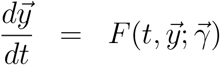

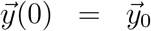

where 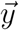 is a vector of state variables, 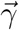 is the vector of parameters, and 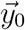 are the initial conditions of the system.
- Quantities of Interest (QoIs; *Q_i_*) that are functions of the state variables and model parameters,

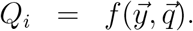

In total, sensitivity analysis assesses and quantifies the relationship between individual input parameters (*q_i_*) and the quantities of interest (QoIs; *Q_i_*). This in turn allows one to identify influential parameters and how input perturbations affect output uncertainty.

There are a variety of sensitivity measures that have been introduced. One measure is local sensitivity, which yields a linear relationship between a parameter, *γ_i_*, and a QoI, *Q_j_*, via the derivative, 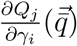. This type of sensitivity analysis is referred to as ‘local’ in the sense that one parameter is varied at a time. Other widely used ‘global’ measures (which simultaneously vary multiple parameters) include Partial Rank Correlation Coefficient, Sobol’ measures, importance measures and screening methods (Sobol, 2001; Marino et al., 2008; Salelli et al., 2004; Saltelli et al., 2019; Jansen, 1999; Jarrett et al., 2017a; Hanthanan Arachchilage and Hussaini, 2021). Global approaches rely on quantifying the relationship between parameter and the QoI based on statistical relationships. They tend to be computationally expensive which is a major difficulty, especially as the number of evaluations increases with the dimensionality of the problem of interest. However, the benefit of global methods is that they can rank the parameters in terms of the magnitudes of their effects while varying all parameters simultaneously. This lowers the chance that the estimated sensitivity indices miss specific contours of the QoI as the parameter landscape changes.

We note that all sensitivity measures depend on where in parameter space we are evaluating the model. Therefore, any rankings of the parameters in terms of the magnitudes of their effects on quantities of interest depends on the location in parameter space. Additionally, local analysis almost always misses important aspects of sensitivity (Saltelli et al., 2019) and a ‘global’ method, where all parameters are varied simultaneous, is preferred (Salelli et al., 2004; Saltelli et al., 2019).

We use Partial Rank Correlation Coefficients (PRCC) (Marino et al., 2008) to quantify the sensitivity analysis. We use PRCC because relationships between parameters and QoIs can be nonlinear and PRCC extends the method to nonlinear relations between parameters and QoIs by focusing on ranked-transformed data. Specifically, PRCC uses rank transformation to transform (potentially nonlinear) monotonic relationships into linear relationships. In the context of noisy observations, where the relationship is not strictly monotonic, more robust conclusions can be made for relationships whose rank transform is approximately linear.

In this manuscript, we use global sensitivity analysis to explore how ecological and epidemiological processes shape the temporal dynamics of a two-host, one pathogen community. The correlation between the parameters and the QoI provides insight into the relationship between biological processes (as reflected in parameter values) and specific outcome measures. In particular a strong, positive correlation imply that small increases in the parameter value lead to qualitatively large changes in the QoI. We use this analysis to identify when and why the magnitudes and signs of the effects of specific ecological and epidemiological parameters on disease dynamics change over time. This in turn yields insight into how the corresponding ecological and epidemiological processes shape disease dynamics, and how those effects depend on the roles and identities of host species. In the following, we define the mathematical model, first developed in Searle et al. (2016), and the biological questions we address. We then describe PRCC in some detail. The results section builds intuition using local sensitivities at equilibrium and then analyzes the qualitative and quantitative dynamics of the PRCC indices in multiple scenarios.

## 2 Mathematical Model

The focal SI-type model describes the dynamics of two-host species and an environmentally transmitted pathogen. In the model, infected individuals release infectious propagules (spores) into the environment when they die and susceptible individuals become infected when they come in contact with the spores. The model was used previously (Searle et al., 2016) to model a laboratory system made up of two water flea species (*Daphnia dentifera* and *D. lumholtzi*) and a shared fungal pathogen (*Metschnikowia bicuspidata*). To match the biology of the empirical system, the model assumes individuals cannot recover from infection (meaning infection is always lethal), infected individuals release spores only when they die, and individuals in both species contribute to and are exposed to the same pool of spores because they share the same habitat.

The dynamics of the densities of susceptible (*S_i_*) and infected (*I_i_*) individuals in each population and the density of spores (*P*) are defined by a system of ordinary differential equations:

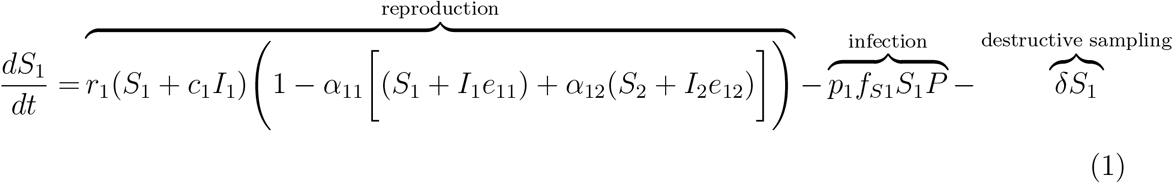

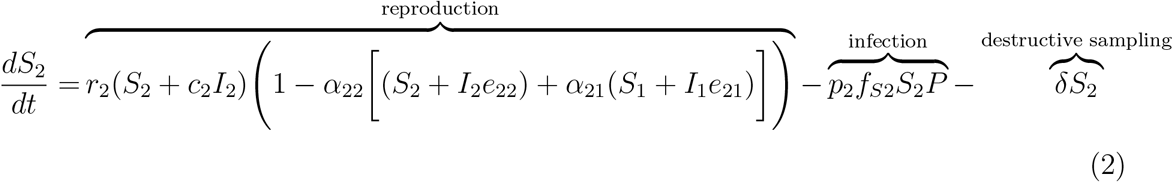

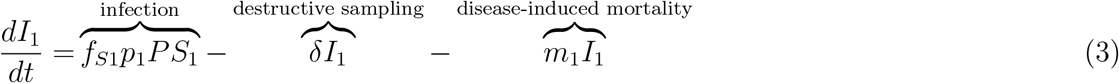

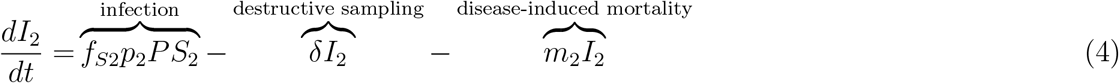

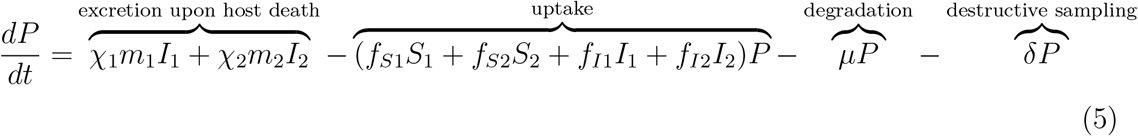

The definitions for all variables and parameters are shown in Table 1. Throughout, we refer to parameters affecting reproduction and competition as ecological parameters (*r_i_*, *c_i_*, *α_ij_*, *e_ij_*, *δ*) and all other parameters as epidemiological parameters (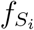, *p_i_*, *m_i_*, *χ_i_*, *μ*). This distinction facilitates the interpretation of the results, but we note that some parameters could be classified in both categories. For example, the filtering rates, 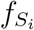 and 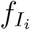, affect both the uptake of infection propagules (an epidemiological process) and uptake of resources (an ecological process).

**Table 1:** Definitions and nominal values of model parameters and variables

| Definition |  | Nominal Value | Units |
| --- | --- | --- | --- |
| <u>Variables</u> |  |  |  |
| $S_i$ | susceptible density for species $i$ | variable | indiv./L |
| $I_i$ | infected density for species $i$ | variable | indiv./L |
| $N_i$ | total density for species $i$ | variable | indiv./L |
| $Y_i$ | proportion infected for species $i$ | variable | unitless |
| $P$ | spore density | variable | spore/L |
| <u>Ecological Parameters</u> |  |  |  |
| $r_1$ | exponential growth rate of host 1 | 0.206 | 1/day |
| $r_2$ | exponential growth rate of host 2 | 0.246 | 1/day |
| $c_1, c_2$ | proportional reduction in growth rate for $I_i$ | 0.75 | unitless |
| $\alpha_{11}$ | intraspecific competition coefficient for host 1 | 1/97.5 | 1/indiv. |
| $\alpha_{22}$ | intraspecific competition coefficient for host 2 | 1/12.8 | 1/indiv. |
| $\alpha_{12}$ | interspecific competition coefficient for effect of host 2 on 1 | 2.63 | unitless |
| $\alpha_{21}$ | interspecific competition coefficient for effect of host 1 on 2 | 0.01 | unitless |
| $e_{ij}$ | reduction in competitive ability of infected individuals | 1 | unitless |
| $\delta$ | average removal rate due to destructive sampling | 0.013 | 1/day |
| <u>Epidemiological Parameters</u> |  |  |  |
| $p_1$ | per spore probability of infection for host 1 | $1.45 \times 10^{-5}$ | 1/spore |
| $p_2$ | per spore probability of infection for host 2 | $4.87 \times 10^{-5}$ | 1/spore |
| $f_{S1}$ | filtering rate for susceptible individuals of host 1 | 0.0348 | 1/L/day |
| $f_{S2}$ | filtering rate for susceptible individuals of host 2 | 0.0361 | 1/L/day |
| $f_{I1}$ | filtering rate for infected individuals of host 1 | 0.0186 | 1/L/day |
| $f_{I2}$ | filtering rate for infected individuals of host 2 | 0.0171 | 1/L/day |
| $m_1, m_2$ | disease induced mortality rate | 0.05 | 1/day |
| $\chi_1$ | spore burst size for host 1 | 120000 | spore/indiv. |
| $\chi_2$ | spore burst size for host 2 | 124000 | spore/indiv. |
| $\mu$ | spore degradation rate | 0.5 | 1/day |

All parameter estimates are taken from Searle et al. (2016).

The terms in equations 1 and 2 define for each species (in order) the total reproduction rates, infection rates, and rates of mortality due to destructive sampling. For host species *i*, *r_i_* and *r_i_c_i_* are the maximum exponential growth rates of susceptible and infected individuals, respectively; *α_ii_* is the intraspecific competition parameter for susceptible individuals; *α_ij_* is the relative effect of interspecific competition of species *j* on species *i*; and *e_ij_* is the competitive effect of infected individuals of species *j* relative to the competitive effect of susceptible individuals of species *j*. When written in the traditional Lotka-Volterra form, the intraspecific competition coefficient for susceptible individuals for host *i* is *α_ii_*, the intraspecific competition coefficient for infected individuals of host *i* is *α_ii_e_ii_*, and the analogous interspecific competition coefficients are *α_ii_α_ij_* and *α_ii_α_ij_e_ij_*. The infection rates are the product of the per spore probability of infection (*p_i_*), the filtering rate of susceptible individuals 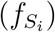 and the densities of susceptible individuals and spores.

The terms in equations 3 and 4 define for each species, the rates of infection 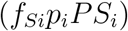 and mortality due to destructive sampling (*δI_i_*) and infection (*m_i_I_i_*). In equation (5), the terms define rates of release of spores by infected individuals when they die (*χ_i_m_i_I_i_*), uptake of spores by susceptible 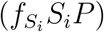 and infected 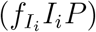 individuals in each population, and loss of spores due to degradation (−*μP*) and destructive sampling (*δP*).

The quantities of interest (QoIs) in this study are the proportion of infected individuals in each population (i.e., *infection prevalence*), defined by *Y_i_* = *I_i_*/*N_i_* where the total density of each population is *N_i_* = *S_i_*+*I_i_*. It is useful to write down the system of equations for the dynamics of the total density and infection prevalence for each population,

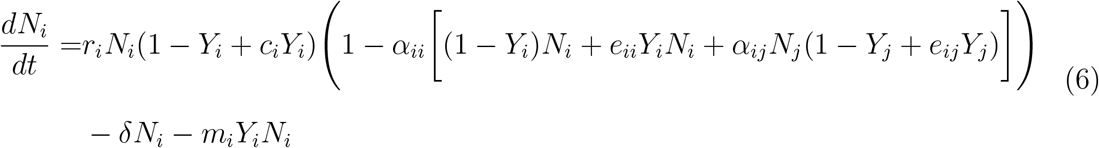

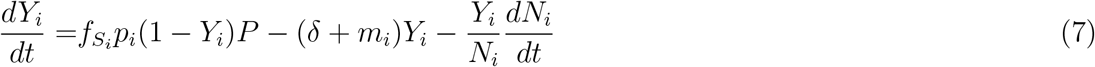

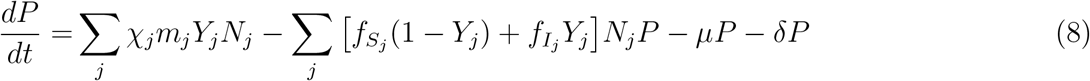

The terms in equation (6) define how the total density of each population changes due to reproduction and mortality. The terms in equation (7) define how the disease prevalence increases due to transmission, decreases due to mortality, and changes as the density of susceptible individuals increases or decreases. Equation (8) is identical to equation (5).

The nominal values for all parameters are shown in Table 1 and were taken from Searle et al. (2016). Briefly, the parameters were estimated in Searle et al. (2016) in the following way. The epidemiological parameter values (e.g., probabilities of infection, spore burst sizes, and mortality rates) were estimated using individual-level experiments where individual *D. dentifera* were exposed to fungal spores and disease-related characteristics were measured. The ecological parameter values (exponential growth rates and competition coefficients) were estimated by fitting equations (1) and (2) to time series data where one or both host species were present and the pathogen was absent. The value for the spore degradation rate (*μ* = 0.5) was taken from a range of values (0 ≤ *μ* ≤ 0.75) within which there was qualitative agreement between the model predictions and experiments where both host species were present with the pathogen.

The models have a unique coexistence equilibrium where all densities are positive. We refer to this equilibrium as the endemic equilibrium and denote its densities using 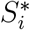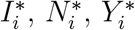, and *P*\*.

## 3 Problem Statement and cases

The goal of this study is to use sensitivity analysis to explore how ecological and epidemiological processes shape the temporal dynamics of disease prevalence in model (1)-(5). To do this, we focus on answering the following questions.

1. How do the magnitudes and signs of the sensitivities of host prevalence to model parameters change over time?
2. When the pathogen is introduced into a disease-free two-host community, how do the temporal patterns depend on the identities of the host species?
3. When a second host species is introduced into a single-host community with an endemic pathogen, how do the temporal patterns depend on which host species is the resident and which is the invader?

We answer these questions by applying sensitivity analysis to simulations of the model in three cases:

Case 1: A moderate amount of spores are added to a disease-free two-host community at equilibrium.
Case 2: Host 2 is introduced at low densities to the single-host endemic equilibrium for host 1. Here, host 1 is the resident and host 2 is invader.
Case 3: Host 1 is introduced at low densities to the single-host endemic equilibrium for host 2. Here, host 2 is the resident and host 1 is invader.

Case 1 identifies if and how the specific characteristics of each host species affect disease dynamics when the pathogen is introduced into the community. Comparing Cases 2 and 3 identifies how the identities of the resident and invader hosts influence disease dynamics.

Specific details about the simulations are the following. In case 1, the initial conditions for the simulations are

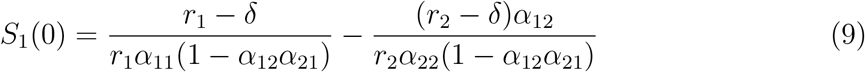

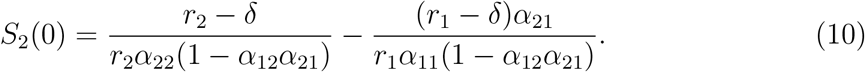

with *I*_1_(0) = 0, *I*_2_(0) = 0, and *P*(0) = 25000. In cases 2 and 3, the system is initiated with nonzero densities for the resident host and spores and the model is run for 2000 days to ensure convergence to the single-host endemic equilibrium. Day 0 is defined as the day the invading host is introduced. The invading host is introduced at 1% of the susceptible density of the resident host (e.g., if host 2 is the invader, then *S*_2_(0) = 0.01 × *S*_1_(0)).

## 4 Global Sensitivity Analysis using PRCC

Our metric for computing global sensitivities is Partial Rank Correlation Coefficients (PRCC) (Marino et al., 2008; Hanthanan Arachchilage and Hussaini, 2021; Wentworth et al., 2016; Jarrett et al., 2017b; Hamby, 1995; Blower and Dowlatabadi, 1994). PRCC quantifies how the variations in the model parameters affect the values of the quantities of interest (QoIs). The method involves sampling parameter space, rank-transforming both the input parameter samples and the computed values for the QoI, and using multiple linear regression to remove the effects from other parameters.

First, PRCC begins with *N* points sampled from the *M*-dimensional parameter space, 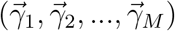, and *N* values of the quantity of interest evaluated at those parameter values, (*Q*_1_, *Q*_2_, …, *Q_N_*). This yields a set of *N* points in *M* + 1 dimensional space. There are a variety of methods that can be used to sample the *M*-dimensional parameter space, including low discrepancy Sobol’ sequences, quasi-Monte Carlo samples and random sampling. In this study, we sample parameter space using Latin Hypercube Sampling (LHS), which partitions each parameter into equally probable intervals so that the hypercube of parameter space is divided into equally probably cubes. Then each parameter is sampled without replacement, to generate *N* random parameter sets. Figure 1 provides a schematic representation of the Latin Hypercubic sampling approach for a Gaussian distribution. Here, five equally probable sample space partitions are generated by dividing the cumulative distribution function (i.e., CDF). Then equal number of samples can be pulled from each partition to create a low discrepancy Gaussian distribution. With no prior knowledge of the variation in parameters, it is typical to assume a uniform distribution which reduces inferred bias. In this study, we assume that the parameter space of each individual input parameter is uniformly distributed with upper and lower bounds of ±10% of its nominal parameter value.

**Figure 1:**
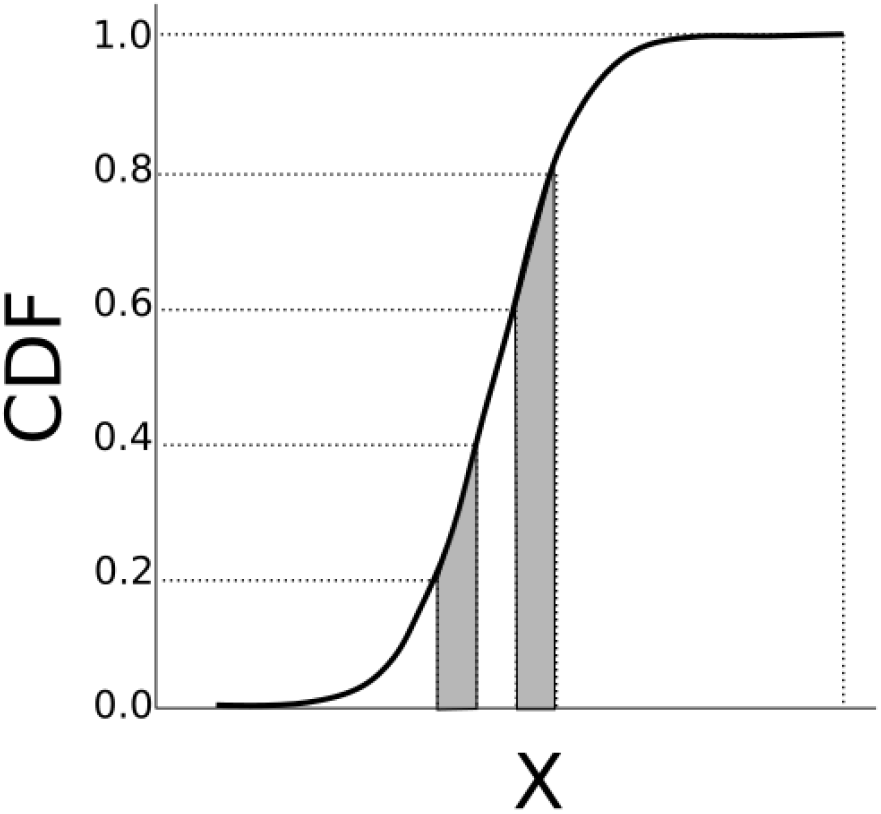
Schematic of the Latin Hypercube sampling showing equiprobable partitioning of the sample space, here obtained from a Gaussian distribution. The x-axis is the sample space and it is equally probable to pull a sample from each of the grey and white bins.

Second, to minimize the effects of nonlinearities, the QoI values and the sampled parameter values are rank-transformed and all calculations are performed on the rank values, rather than the values themselves (see Figure 2); this yields Partial Rank Correlation Coefficients (PRCC). Rank transformation is needed because sensitivity indices such as Pearson correlation coefficients require the variation between the input parameters and a QoI to be linear in order to obtain an accurate estimation (Wentworth et al., 2016; Marino et al., 2008). However, linear relationships seldom arise in real life applications. The rank transformation addresses this issue by ordering the sampled parameters values from most negative to most positive and then assigning an integer (in order) to each sampled parameter value. The same transformation is done to the QoI values. This transformation linearizes the data; see Figure 2c,d for an illustration.

**Figure 2:**
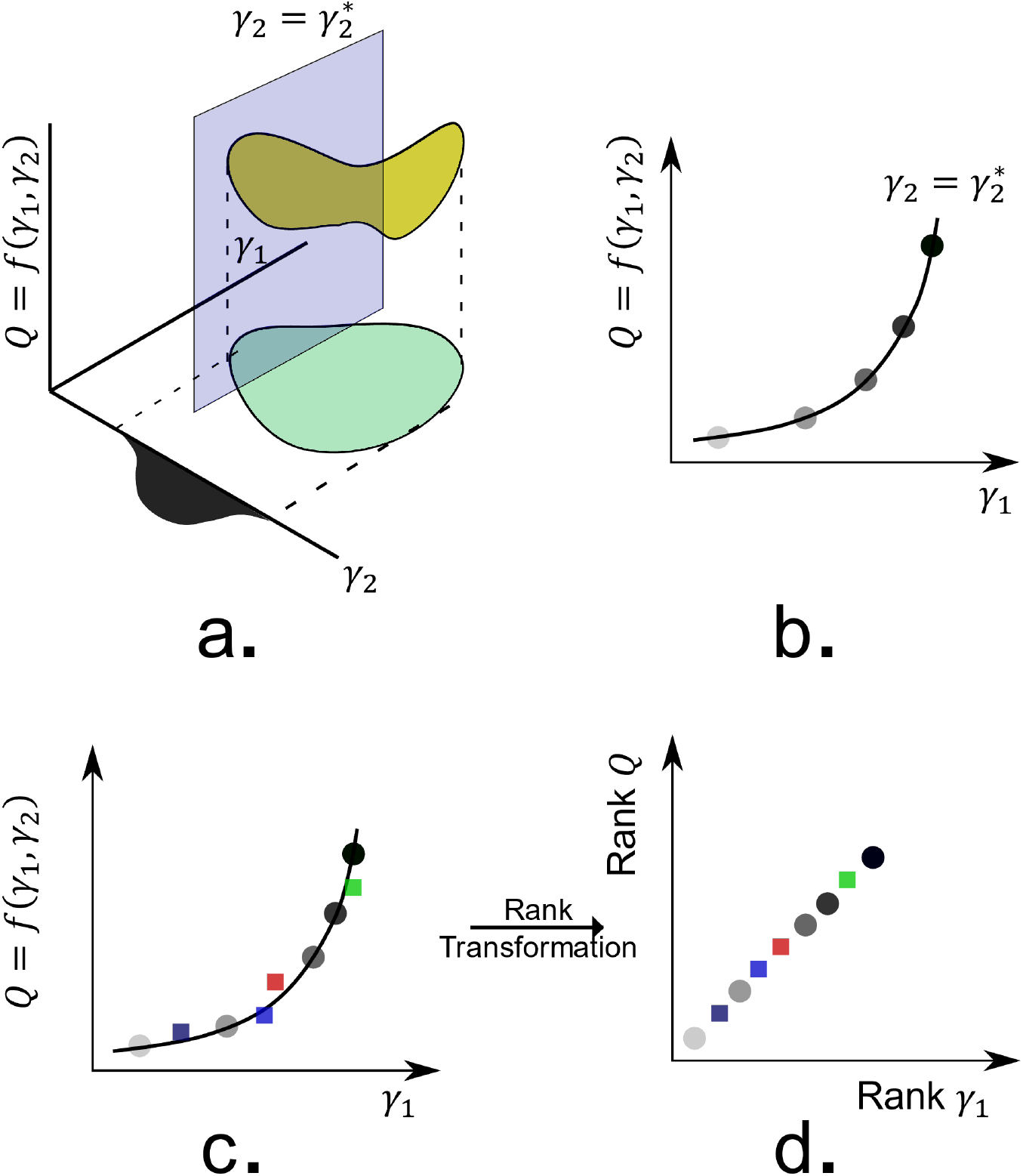
Schematic for computing PRCC. (a) Values of the quantity of interest (*Q*, yellow) for a region of parameter space (*γ*_1_ and *γ*_2_, green). The probability density function for the parameter *γ*_2_ is shown in black on the *γ*_2_-axis. (b) Variation in the quantity of interest in the *γ*_1_ direction at a nominal value 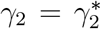. Gray scale circles indicate values of the quantity of interest at specific values of *γ*_1_ when 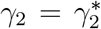. (c) Variation in the quantity of interest in the *γ*_1_ direction when parameters are sampled over the joint distribution of points in *γ*_1_*γ*_2_-parameter space. Gray scale circles correspond to points in panel (b) and colored squares scattered around the curve correspond to points where 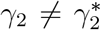. (d) Rank transformation yields a linear relationship between the rank transformed quantity of interest, Rank *Q*, and the rank transformed parameter, Rank *γ*_1_.

Importantly, in order for linearization using rank-transformation to be successful, the relationship between the input parameters and the QoI needs to be monotonic (Marino et al., 2008). For all parameters considered in this study, there was a monotonic relationship between each parameter and each QoI. Monotonicity for each parameter value, *q_i_*, was verified by freezing all variables except for *q_i_* at their nominal values and varying parameter *q_i_* within its allowed range. In Figures 2.a and 2.b we show a schematic for a QoI of the form *Q* = *f*(*γ*_1_, *γ*_2_). Because the relationship between the QoI (*Q*) and the parameter *γ*_1_ is monotonic (Figure 2.b), the relationship between the rank-transformed QoI and parameter values is linear (Figure 2.d). If the relationship between the QoI and the parameter was not monotonic, then the relationship between the rank-transformed QoI and parameter values could be nonlinear and the estimated PRCC could be a poor representation of the relationship.

We note that PRCC can be interpreted statistically in terms of the correlation between the parameters and the QoI (Marino et al., 2008; Hanthanan Arachchilage and Hussaini, 2021; Salelli et al., 2004). The estimation of PRCC sensitivity indices for a parameter of interest (i.e., *q_i_*) consists of 3 steps. As mentioned earlier, the first step is to rank-transform both sampled input parameters and the estimated QoI values. Let *γR_i_* be the rank-transformed parameters corresponding to parameter *γ_i_* and *QR* be the rank-transformed QoI values. Secondly, we remove the influence of other parameters (*γR*_≠*i*_) by estimating the influence of *γR*_≠*i*_ on *γR_i_* and *QR* respectively using multiple linear regression and subtracting from the original value (i.e., *γR_i_* and *QR*). Specifically, the respective predictions for the ranked parameter *γ_i_* and the ranked QoI are

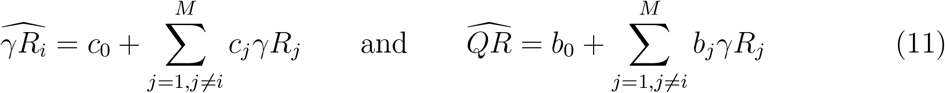

where *c_o_, c_j_, b*_0_ and *b_j_* are the inferred regression coefficients. Once 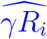 and 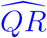 are calculated, the corresponding residues (i.e., 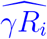 and 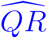) are calculated by subtracting from the original values to yield,

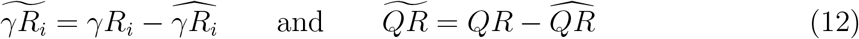

Thirdly, the PRCC sensitivity index for parameter *γ_i_* can be calculated as the correlation coefficient between the 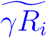 and 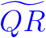,

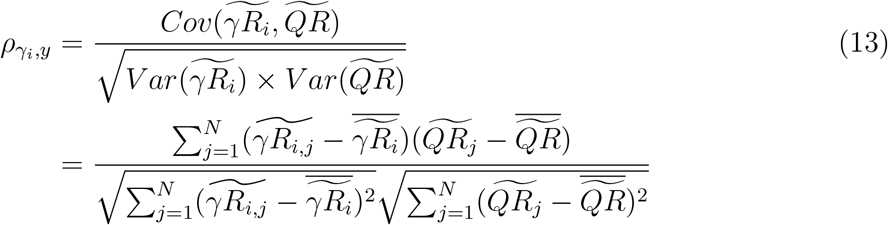

where overbars denote the means. Positive and negative values imply positive and negatively correlations, respectively, between the rank-transformed parameter and rank-transformed QoI. Higher absolute values indicate a higher correlation between the rank-transformed parameter and rank-transformed QoI.

## 5 Results

The results are organized as follows. Section 5.1 builds intuition about the expected signs of the PRCC sensitivity indices using the differential equations in the model (section 5.1.1) and local sensitivities evaluated at the two-host endemic equilibrium (section 5.1.2). Section 5.2 compares those expected signs with the signs of the PRCC sensitivities computed for Cases 1-3 and focuses on explaining when and why the signs of the global sensitivities agree and disagree with expected signs. Section 5.3 focuses on the magnitudes of the PRCC sensitivities. That section explores the quantitative dynamics of the PRCC sensitivities and explains the biological insight that is gained by analyzing the temporal dynamics in Case 1 (section 5.3.1) and by comparing temporal dynamics of Cases 2 and 3 (section 5.3.2).

### 5.1 Building intuition about the expected signs of the global sensitivities

#### 5.1.1 Expected signs based on model equations

One way to build intuition about the signs of the global sensitivities is to compute the partial derivatives of the growth rates for disease prevalence (*dY_i_*/*dt*) and spore density (*dP*/*dt*) with respect to the model parameters. The mathematical reasoning for considering how parameter values affect the derivative of a state variable is the following. For one-dimensional systems, changes in parameter values have effects of the same sign on the state variable and its derivative at all points in time and at equilibrium. Specifically, consider the non-autonomous ODE *dQ/dt* = *F* (*Q, γ*) with parameter *γ* and initial condition *Q*(0) = *Q*_0_. Let 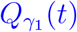 denote the solution for a particular parameter value *γ_i_*. If *F* is an increasing function of *γ*, i.e., *∂F/∂γ*> 0, then 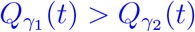 for all *γ*_1_ > *γ*_2_ for all time and if *F* is a decreasing function of *γ*, i.e., *∂F/∂γ*< 0, then 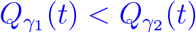 for all *γ*_1_ > *γ*_2_ for all time. In addition, at a stable equilibrium, *Q^*^*, of the ODE, which must satisfy *∂F/∂Q*|_*Q*\*_< 0, we get that *∂Q^*^/∂γ*|_*N*\*_ = −(*∂F/∂γ*)/(*∂F/∂Q*)|_*Q*\*_, which has the same sign as *∂F/∂γ*. Thus, changes in parameter values that increase or decrease the derivative of a state variable also increase or decrease, respectively, the value of the state variable at all points in time and at equilibrium. For higher dimension systems, changes in parameter values have effects of the same sign on the state variable and its derivative over sufficiently short time scales. However, over longer time scales and at equilibrium, the effects can be of opposite signs due to the feedbacks between all state variables. This means that the effects of changes in parameter values on the derivative of a state variable are likely to yield insight into how changes in parameter values affect the value of a state variable over short time scales and they can, but do not always, yield insight into how changes in parameter values affect the value of a state variable over long time scales.

Based on the above, the intuition is that disease prevalence in host *i* will increase and decrease with variation in a parameter based on whether the parameter causes the growth rates of disease prevalence and spore density to increase or decrease, respectively.

In particular, one might expect that prevalence in host *i* (*Y_i_* = *I_i_*/*N_i_*) and the parameter *γ* are (a) positive correlated if the partial derivatives of *dY_i_*/*dt* or *dP*/*dt* with respect to *γ* are positive (i.e, [*∂*/*∂γ*][*dY_i_*/*dt*] > 0 or [*∂*/*∂γ*][*dP*/*dt*] > 0), (b) negatively correlated if the partial derivatives of *dY_i_*/*dt* or *dP*/*dt* with respect to *γ* are negative (i.e, [*∂*/*∂γ*][*dY*_1_/*dt*] < 0 or [*∂*/*∂γ*][*dP*/*dt*] < 0), and (c) uncorrelated if *dY_i_*/*dt* and *dP*/*dt* do not depend on *γ* (i.e, [*∂*/*∂γ*][*dY*_1_/*dt*] = 0 and [*∂*/*∂γ*][*dP*/*dt*] = 0).

The signs of the derivatives of the growth rates are summarized in the columns labeled “Model Eqns.” in Table 2. We note two issues with this intuition gained from this approach. First, in general, this approach is unlikely to predict the signs of the global sensitivities in all cases because it ignores feedbacks between the variables. For example, *dY_i_*/*dt* does not depend on probability of infection for the other host species (*p_j_*, *i* ≠ *j*), but the value of *p_j_* will have an indirect effect on the magnitude of *dY_i_*/*dt* because the differential equations are coupled. Second, for some parameters the signs of the derivatives of the *dY_i_*/*dt* and *dP*/*dt* equations differ, which makes it difficult to predict the sign of the correlation between prevalence and the parameter. For example, *dY*_1_/*dt* is an increasing function of the host filtering rate 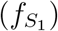 whereas *dP*/*dt* is a decreasing function.

**Table 2:** Computed signs of global and local sensitivities

| Parameter | Quantity of Interest 1, $Y_1 = I_1/N_1$ | | | | | Quantity of Interest 2, $Y_2 = I_2/N_2$ | | | | |
| --- | --- | --- | --- | --- | --- | --- | --- | --- | --- | --- |
| | Model<br>Eqns.* | Case 1<br>$t > 0$ | Case 2<br>$t > 0$ | Case 3<br>$t > 0$ | LSA<br>Eq.* | Model<br>Eqns.* | Case 1<br>$t > 0$ | Case 2<br>$t > 0$ | Case 3<br>$t > 0$ | LSA<br>Eq.* |
| <u>Ecological</u> |  |  |  |  |  |  |  |  |  |  |
| $r_1$ | — | $-\uparrow+$ | $+\updownarrow+$ | $-\updownarrow+$ | + | | $-\uparrow+$ | $+\updownarrow+$ | $+\updownarrow+$ | + |
| $r_2$ | | $+\uparrow+$ | $+\updownarrow+$ | $+\updownarrow+$ | + | — | $+\uparrow+$ | $-\updownarrow+$ | $+\updownarrow+$ | + |
| $C_1$ | — | $-\updownarrow+$ | $+\updownarrow+$ | $-\updownarrow+$ | + | — | $-\updownarrow+$ | $+\updownarrow+$ | $+\updownarrow+$ | + |
| $C_2$ | | $+\uparrow+$ | $-\uparrow+$ | $+\updownarrow+$ | + | | $-\uparrow+$ | $-\updownarrow+$ | $+\updownarrow+$ | + |
| $\alpha_{11}$ | + | $+\downarrow-$ | $-\uparrow-$ | $+\updownarrow-$ | — | | $+\downarrow-$ | $-\updownarrow-$ | $-\updownarrow-$ | — |
| $\alpha_{12}$ | + | $+\downarrow-$ | $+\updownarrow-$ | $+\updownarrow-$ | — | | $+\downarrow-$ | $+\updownarrow-$ | $+\updownarrow-$ | — |
| $\alpha_{21}$ | | $-\updownarrow-$ | $+\downarrow-$ | $-\updownarrow-$ | — | + | $-\updownarrow-$ | $+\updownarrow-$ | $-\updownarrow-$ | — |
| $\alpha_{22}$ | | $-\updownarrow-$ | $-\downarrow-$ | $-\updownarrow-$ | — | + | $-\updownarrow-$ | $+\updownarrow-$ | $-\updownarrow-$ | — |
| $e_{ij}$ | + | $+\downarrow-$ | $-\updownarrow-$ | $-\updownarrow-$ | — | + | $+\downarrow-$ | $-\updownarrow-$ | $-\uparrow-$ | — |
| $\delta$ | — | $-\downarrow-$ | $-\downarrow-$ | $-\updownarrow-$ | — | — | $-\downarrow-$ | $-\downarrow-$ | $-\updownarrow-$ | — |
| <u>Epidemiological</u> |  |  |  |  |  |  |  |  |  |  |
| $p_1$ | + | $+\updownarrow+$ | $+\updownarrow+$ | $+\uparrow+$ | + | | $+\uparrow+$ | $+\downarrow+$ | $+\uparrow+$ | + |
| $p_2$ | | $+\updownarrow+$ | $-\uparrow+$ | $+\downarrow+$ | + | + | $+\downarrow+$ | $+\uparrow+$ | $+\downarrow+$ | + |
| $f_{S_1}$ | + | $+\updownarrow+$ | $+\uparrow+$ | $+\updownarrow+$ | + | — | $-\uparrow+$ | $+\updownarrow+$ | $+\updownarrow+$ | + |
| $f_{S_2}$ | — | $-\updownarrow+$ | $-\updownarrow+$ | $+\updownarrow+$ | + | + | $+\downarrow+$ | $+\uparrow+$ | $+\downarrow+$ | + |
| $f_{I_1}$ | — | $-\downarrow-$ | $-\downarrow-$ | $+\downarrow-$ | — | — | $+\downarrow-$ | $-\downarrow-$ | $+\downarrow-$ | — |
| $f_{I_2}$ | — | $-\downarrow-$ | $-\downarrow-$ | $-\uparrow-$ | — | — | $-\downarrow-$ | $-\downarrow-$ | $-\uparrow-$ | — |
| $m_1$ | + | $+\updownarrow+$ | $+\downarrow+$ | $-\updownarrow+$ | + | + | $+\uparrow+$ | $+\updownarrow+$ | $+\uparrow+$ | + |
| $m_2$ | + | $+\updownarrow+$ | $-\uparrow+$ | $+\downarrow+$ | + | + | $-\updownarrow-$ | $-\downarrow-$ | $+\updownarrow-$ | — |
| $\chi_1$ | + | $+\uparrow+$ | $+\downarrow+$ | $+\uparrow+$ | + | + | $+\uparrow+$ | $+\downarrow+$ | $+\uparrow+$ | + |
| $\chi_2$ | + | $+\updownarrow+$ | $+\uparrow+$ | $+\downarrow+$ | + | + | $+\updownarrow+$ | $+\uparrow+$ | $+\downarrow+$ | + |
| $\mu$ | — | $-\downarrow-$ | $-\downarrow-$ | $-\uparrow-$ | — | — | $-\downarrow-$ | $-\downarrow-$ | $-\uparrow-$ | — |

Plus (+) and minus (−) signs mean positive and negative values, respectively. The ordering of the signs indicates how the sign of the global sensitivity changes as time increases from zero (left sign) to infinity (right sign). Up arrows (↑) and down arrows (↓) mean that the global sensitivity value changes monotonically as time increases. Up-down arrows (↑↓) mean that the global sensitivity value changes in a non-monotonic way.
\* Signs based on the derivatives of the model equations (Model Eqns.) and the local sensitivities evaluated at the two-host endemic equilibrium (LSA Eq.); see text for details. Plus (+) and minus (−) signs mean positive and negative values, respectively. Blank entries mean that the $dY_i/dt$ and $dP/dt$ equations are independent of the parameter.

#### 5.1.2 Signs of local sensitivities computed at equilibrium

Another way to build intuition about the signs of global sensitivities is to compute local sensitivities at equilibrium. While global sensitivities are often better to use than local sensitivities (as discussed in the Introduction), we compute the local sensitivities at equilibrium for three reasons. First, for the nominal parameter values, the simulations for Cases 1-3 converge to the equilibrium used to compute the local sensitivities. Second, the local sensitivity of the equilibrium value of a QoI, *Q^*^*, to a parameter, *γ*, can be computed analytically via the partial derivatives *∂Q^*^/∂γ*; see appendix A.1 for details. In addition, Cortez and Duffy (2021) and Cortez (2021) analyzed generalized versions of models (1)-5) and (6)-(7) and showed that the analytical formulas can be interpreted in terms of indirect feedbacks between variables. Thus, the analytically calculated local sensitivities at equilibrium, allow us to connect biological mechanism with computed signs of the local sensitivities. Three, when the parameters are set to the nominal values, the simulations for Cases 1, 2, and 3 converge to the (unique) endemic equilibrium used to compute the local sensitivities. Combined, this suggests that insight gained about the mechanisms determining the signs of the local sensitivities at equilibrium may also yield insight into the mechanisms determining the signs of the global sensitivities at sufficiently large time scales. Moreover, if the signs of the local sensitivities at equilibrium and the signs of the global sensitivities at shorter time scales agree, then the mechanistic insight gained from the local sensitivities provides a good starting point for understanding the mechanisms driving the signs of the global sensitivities at short time scales. In total, studying the local sensitivities at equilibrium is useful because they provide a good starting point when trying to connect biological mechanism with observed patterns in the global sensitivities.

The signs of the local sensitivities for the nominal parameter values are given in the columns labeled “Eq. Local Sensitivity” in Table 2; see appendix A.2 for details about each individual parameter. The overall patterns are that infectious prevalence in either host increases with higher host reproduction rates (larger *r_i_* and *c_i_* and smaller *α_ij_* and *e_ij_*), higher rates of infection (larger *p_i_* and 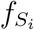), higher spore release rates (larger *χ_i_* and *m_i_*), and lower loss rates of spores (smaller 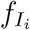, *μ*, and *δ*). The intuition is that higher host densities, higher transmission rates, and greater spore density lead to greater contact between susceptible individuals and spores, which leads to more infections and higher prevalence. The only exception is that the local sensitivity for infection prevalence in host 2 (*I*_2_/*N*_2_) is negative for its own mortality rate (*m*_2_). The reason is that host 2 density is very low and while increased mortality leads to greater release rates of spores, it also reduces the population size of the second host, which leads to fewer infected individuals of host 2.

#### 5.1.3 Comparisons and limitations of predicted signs

The intuition gained from the previous two approaches agrees for some parameters and disagrees for others; see appendix A.3 for discussions about why disagreement occurs for specific parameters. Agreement and disagreement both provide useful biological insight because they identify ecological and epidemiological processes whose influence on disease prevalence may or may not vary over time. Cases of agreement could suggest that the signs of the global sensitivities are the same at all points in time, which would mean that the corresponding biological processes have effects of the same sign on infection prevalence at all time scales. Applying this logic to Table 2, we predict that infection probabilities (*p_i_*) and spore burst sizes (*χ_i_*) have positive effects at all time scales and filtering by infected individuals 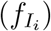, spore degradation (*μ*), and destructive sampling (*δ*) have negative effects at all time scales. In comparison, cases of disagreement suggest that the signs of the global sensitivities are likely to differ over time, which would mean that the corresponding biological processes can have effects of different signs on infection prevalence at different points in time. Applying this to Table 2, we predict that host reproduction (*r_i_*, *c_i_*), host competition (*α_ij_*, *e_ij_*), host mortality (*m_i_*), and filtering by susceptible individuals 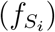 could have positive and negative effects on infection prevalence.

We note three things about the predictions. First, in general, the predictions from the two approaches differ because the local sensitivities account for all direct and indirect effects between the model variables whereas the derivatives of the model equations only account for direct effects. Second, as explained above, the predictions from each approach have the potential to identify the mechanisms determining the signs of the global sensitivities. However, an important limitation is that local sensitivities at equilibrium and the derivatives of the model equations do not account for variation in the initial state of the system, i.e., the initial conditions of the model. In particular, the initial conditions in Cases 1-3 are defined by equilibria of the system, whose locations in phase space depend on the model parameters (e.g., the location of the disease-free two-host equilibrium depends on the competition coefficients *α_ij_*). Thus, the signs and magnitudes of the global sensitivities could be influenced by the initial conditions of the system. Third, while local sensitivity accounts for feedbacks between variables, it only provides a limited exploration of parameter space and phase space. Specifically, the local sensitivity calculations only compute infinitesimal variations in one parameter at a time (and implicitly assumes the gradients stay the same throughout the entire parameter space), which provides a very limited exploration of parameter space, and they are only evaluated at the endemic equilibrium, which provides a very limited exploration of phase space. Consequently, while the local sensitivities have the potential to explain some mechanisms determining the signs of the global sensitivities (as explained above), those predictions may be inaccurate as parameters are varied over large ranges and state variables vary over time.

### 5.2 Analysis of temporal changes in the signs of global sensitivities

We now analyze the temporal changes in the signs of the global sensitivities for disease prevalence in each host species. We focus on parameters whose global sensitivities differ in sign at some time point from the local sensitivities at equilibrium. This helps identify the general mechanisms explaining why the effects of specific ecological and epidemiological processes on disease prevalence can differ in sign over different time scales. We refer to parameters whose PRCC values exceed 0.6 for at least one host at some point in time (i.e, *max*|*PRCC*(*t*)| ≥ 0.6) as the most sensitive parameters. To simplify the presentation in the main text, we primarily discuss parameters whose PRCC values exceed 0.75 and refer the reader to appendix B for discussion about the other parameters whose PRCC values exceed 0.6. The temporal dynamics for the most sensitive parameters are plotted in Figures 3-5. Table 2 roughly summarizes the trends. In particular, for each entry in Table 2, the first symbol is the sign at *t* ≈ 0, the third symbol is the sign at *t* = 3000, and the middle symbol shows whether the PRCC value increases monotonically (up arrow), decreases monotonically (down arrow) or changes in a non-monotonic way (up-down arrow).

**Figure 3:**
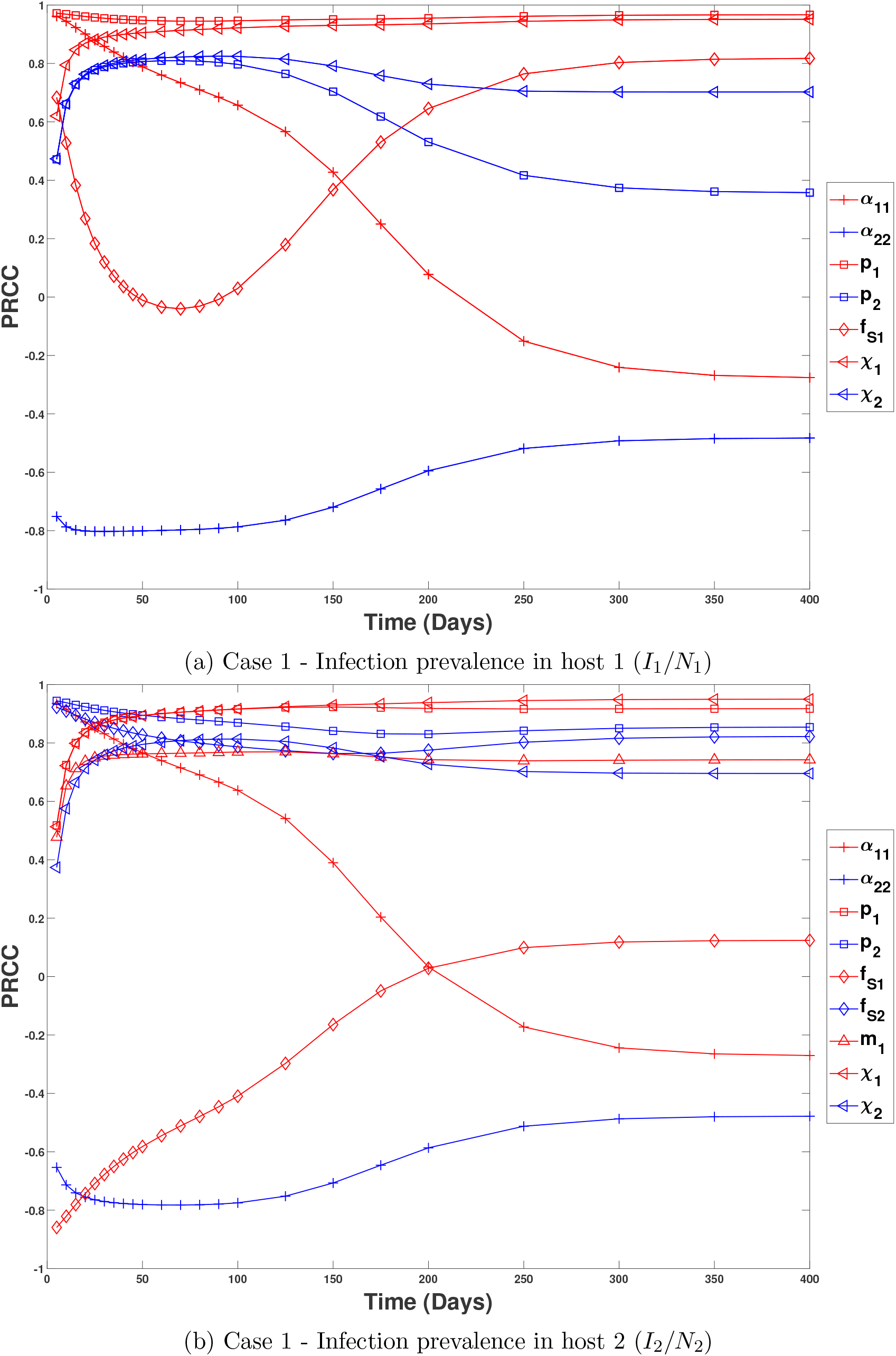
Temporal patterns in the global sensitivities for Case 1. In Case 1, the pathogen is introduced into the two-host community at equilibrium. Each panel shows the global sensitivities for parameters satisfying max(|*PRCC*(*t*)| ≥ 0.75 for (top) disease prevalence in host 1, *I*_1_/*N*_1_, and (bottom) disease prevalence in host 2, *I*_2_/*N*_2_. The specific parameter associated with each curve is given in the legend; red corresponds to parameter values for host 1 and blue corresponds to parameter values for host 2.

**Figure 4:**
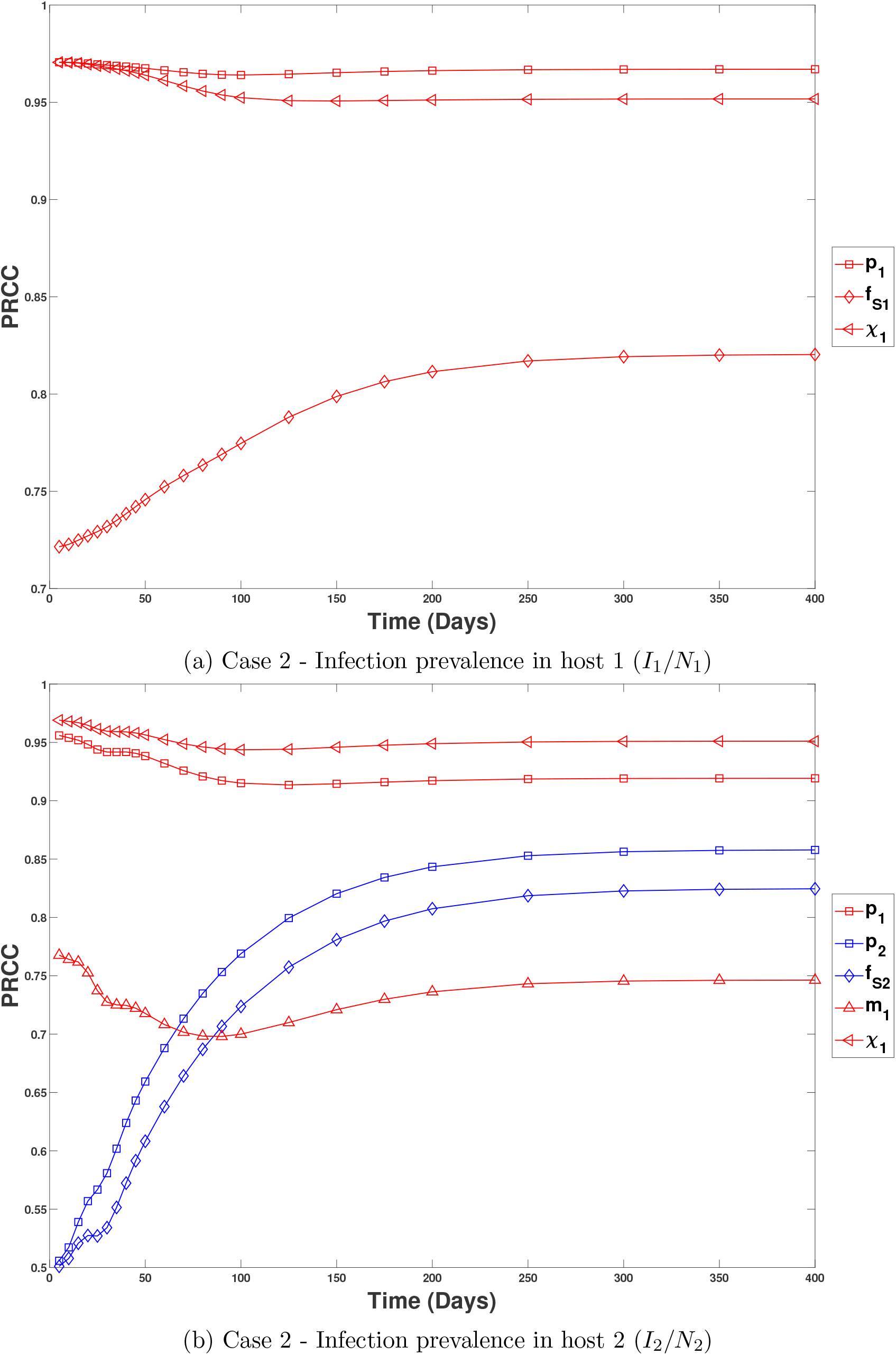
Temporal patterns in the global sensitivities for Case 2. In Case 2, host 2 is introduced into the endemic community made up of host 1 and the pathogen. Each panel shows the global sensitivities for parameters satisfying max(|*PRCC*(*t*)| ≥ 0.75 for (top) disease prevalence in host 1, *I*_1_/*N*_1_, and (bottom) disease prevalence in host 2, *I*_2_/*N*_2_. The specific parameter associated with each curve is given in the legend; red corresponds to parameter values for host 1 (the resident), blue corresponds to parameter values for host 2 (the invader), and green corresponds to parameter values for the pathogen.

**Figure 5:**
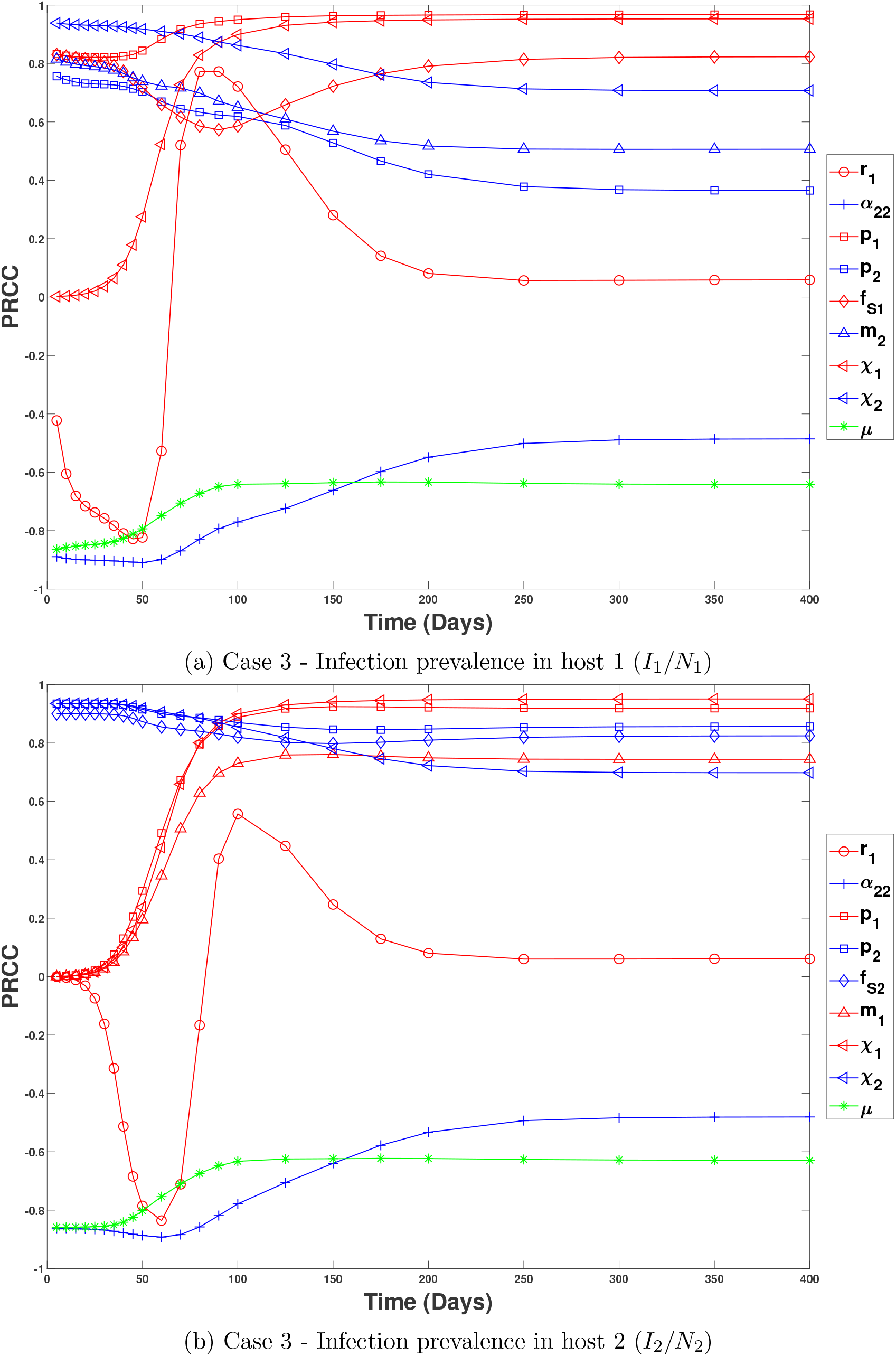
Temporal patterns in the global sensitivities for Case 3. In Case 3, host 1 is introduced into the endemic community made up of host 2 and the pathogen. Each panel shows the global sensitivities for parameters satisfying max(|*PRCC*(*t*)| ≥ 0.75 for (top) disease prevalence in host 1, *I*_1_/*N*_1_, and (bottom) disease prevalence in host 2, *I*_2_/*N*_2_. The specific parameter associated with each curve is given in the legend; red corresponds to parameter values for host 1 (the invader) and blue corresponds to parameter values for host 2 (the resident).

There are three takeaways from Figures 3-5 and Table 2. First, in many instances there is agreement between the signs of the global sensitivities and the signs of the local sensitivities at equilibrium. Specifically, there is agreement (i) for all parameters in all cases at sufficiently large time (i.e., third symbol for each case agrees with the symbol in “LSA Eq.” column); (ii) for each parameter there is agreement for all time points in at least one of the three cases (e.g., signs for *r*_1_ agree for cases 2 and 3, but not case 1); and (iii) for a few parameters (*χ_i_*, *δ*, *μ*) there is agreement for all time points in all three cases. These instances of agreement suggest that, for the range of variation used in the global sensitivity calculations, the signs of the global sensitivities are likely driven by the same mechanisms that explain the signs of the local sensitivities at equilibrium. For example, positive global sensitivities for *χ*_1_ and *χ*_2_ for *δ* and *μ* are likely due to those parameters increasing spore density, which leads to higher contact rates between susceptible individuals and spores and ultimately, more infections and higher infection prevalence. Similarly, negative global sensitivities for *δ* and *μ* are likely due to those parameters decreasing the spore densities, which ultimately leads to fewer infections and lower infection prevalence.

Second, for nearly all of the parameters, there exists at least one case where the signs of the global sensitivities and local sensitivities disagree for finite periods of time. In addition, the signs of the derivatives of the model equations (column “Model Eqns.” in Table 2) and local sensitivities at equilibria (column “LSA Eq.”) do not predict which parameters will have global sensitivities whose signs differ from the signs of the local sensitivities at equilibrium. Thus, the temporal patterns of the global sensitivities show that for almost every ecological or epidemiological process, there exist biologically relevant conditions under which the sign of the effect of that process on infection prevalence in one or both host species changes signs.

The third takeaway is that the specific conditions under which the sign changes occur identify when and how nonlinear interactions between multiple processes lead to unexpected effects of a process on infection prevalence. Our numerical results show that sign changes in the most sensitive parameters occur because of how reproduction and production of spores influences infection prevalence. Brief explanations for the sign changes for the most sensitive parameters in each case are provided below; see appendix B for additional details.

In Case 1, the sensitivities of host 1 prevalence (*I*_1_/*N*_1_) and host 2 prevalence (*I*_2_/*N*_2_) to the intraspecific competition coefficient for host 1 (*α*_11_) change from positive to negative (red plus signs in Figure 3a,b) and the sensitivities of prevalence in both host species to the filtering rate of host 1 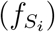 change from positive to negative to positive (red diamonds in Figure 3a,b). The reason for all sign changes is that spore density is initially very low and host 2 produces more spores per infected individual than host 1. Consequently, reduced density and filtering by host 1 results in faster increases in spore density, and thus more infections in both hosts. However, later in time greater density and filtering in host 1 yields higher spore density and more infections in both hosts.

In Case 2 (Figure 4), the signs of the global sensitivities for the most sensitive parameters do not change sign.

In Case 3, the sensitivities of host 1 prevalence (*I*_1_/*N*_1_) and host 2 prevalence (*I*_2_/*N*_2_) to the reproduction rate of host 1 (*r*_1_) exhibit substantial change in both sign and magnitude (red circles in Figure 5a,b). Early in time (day 0 to 75), host 1 is increasing from low densities. Increased reproduction of host 1 causes the density of susceptible individuals of host 1 to increase faster than the density of infected individuals. The sensitivity for host 1 prevalence is negative because faster growth of susceptible individuals than infected individuals necessarily causes host 1 prevalence to decrease. The sensitivity for host 2 prevalence is negative because greater host 1 density results in great uptake of spores, which decreases the infection rate for host 2 and decreases host 2 prevalence. Both global sensitivities are large in magnitude because variation in the growth rate of host 1 has a large effect on how fast host population 1 exponentially grows. Later in time (day 75-150), susceptible density of host 1 overshoots its equilibrium value (see Figure B9), which results in susceptible density of host 1 decreasing over time. This reverses the signs of the global sensitivities because greater reproduction by host 1 means there are more susceptible individuals who can subsequently become infected. As a result, increased reproduction of host 1 yields higher prevalence in host 1, and because higher host 1 prevalence yields greater spore densities, higher prevalence in host 2 as well. Both global sensitivities are large in magnitude because variation in the growth rate of host 1 has a large effect on how the rate of production of susceptible individuals (who can subsequently become infected). As the system converges to equilibrium (day 150 and later), the global sensitivities of both prevalence monotonically decrease to small positive values. The values are small in magnitude because the host growth rates have a small effect on the densities and prevalences at equilibrium. The values are positive because increased reproduction by host 1 leads to greater densities of host 1, which increases host 1 prevalence. In addition, increased host 1 prevalence yields greater spore densities, which results in increased host 2 prevalence.

### 5.3 Comparisons of global sensitivities across cases

Here, we compare the temporal dynamics of the PRCC sensitivity indices across the three cases in order to answer questions two and three posed in Section 3. The first subsection compares the temporal dynamics of the sensitivities for the two host species in Case 1. The second subsection compares the temporal dynamics for the two host species across Cases 2 and 3. We do not compare the temporal dynamics of Case 1 with those of Cases 2 and 3 because the differences in initial conditions make it unclear what biological insight can be gained. We primarily discuss the most sensitivity parameters (defined by max(|*PRCC*(*t*)| ≥ 0.6), with the main text focusing on parameters satisfying max(|*PRCC*(*t*)| ≥ 0.75 (summarized in Table 3) and appendix C focusing on the other parameters whose PRCC values exceed 0.6. Brief explanations are given in each subsection; see appendix C for detailed explanations.

**Table 3:** Comparison of parameters satisfying *max*|*PRCC*(*t*)| ≥ 0.75 in each case

|  | Case 1 |  | Case 2 |  | Case 3 |  |
| --- | --- | --- | --- | --- | --- | --- |
| Parameter | QoI 1 | QoI 2 | QoI 1 | QoI 2 | QoI 1 | QoI 2 |
| | $Y_1 = I_1/N_1$ | $Y_2 = I_2/N_2$ | $Y_1 = I_1/N_1$ | $Y_2 = I_2/N_2$ | $Y_1 = I_1/N_1$ | $Y_2 = I_2/N_2$ |
| $r_1$ | | | | | ✓ | ✓ |
| $\alpha_{11}$ | ✓ | ✓ | | | | |
| $\alpha_{22}$ | ✓ | ✓ | | | ✓ | ✓ |
| $p_1$ | ✓ | ✓ | ✓ | ✓ | ✓ | ✓ |
| $p_2$ | ✓ | ✓ | | ✓ | ✓ | ✓ |
| $f_{S_1}$ | ✓ | ✓ | ✓ | | ✓ | |
| $f_{S_2}$ | | ✓ | | ✓ | | ✓ |
| $m_1$ | | ✓ | | ✓ | | ✓ |
| $m_2$ | | | | | ✓ | |
| $\chi_1$ | ✓ | ✓ | ✓ | ✓ | ✓ | ✓ |
| $\chi_2$ | ✓ | ✓ | | | ✓ | ✓ |
| $\mu$ | | | | | ✓ | ✓ |

To facilitate the comparisons, Figures 6-8 are organized such that such that color denotes species identity for a parameter (blue denotes parameters for host 1 and red denotes parameters for host 2); shape denotes the specific case (circles denote Case 1, squares denote Case 2, and triangles denote case 3); and line type and symbol fill denote whether the species is a resident or an invader (solid curves and filled symbols denote the resident’s parameters whereas dashed curves and open symbols denote the invader’s parameters). Our comparisons focus on identifying when the magnitudes and dynamics of the sensitivities can be predicted by (i) intraspecific versus interspecific processes, (ii) the species’ roles of invader versus resident, or (iii) the species’ identities; see below for details about each.

**Figure 6:**
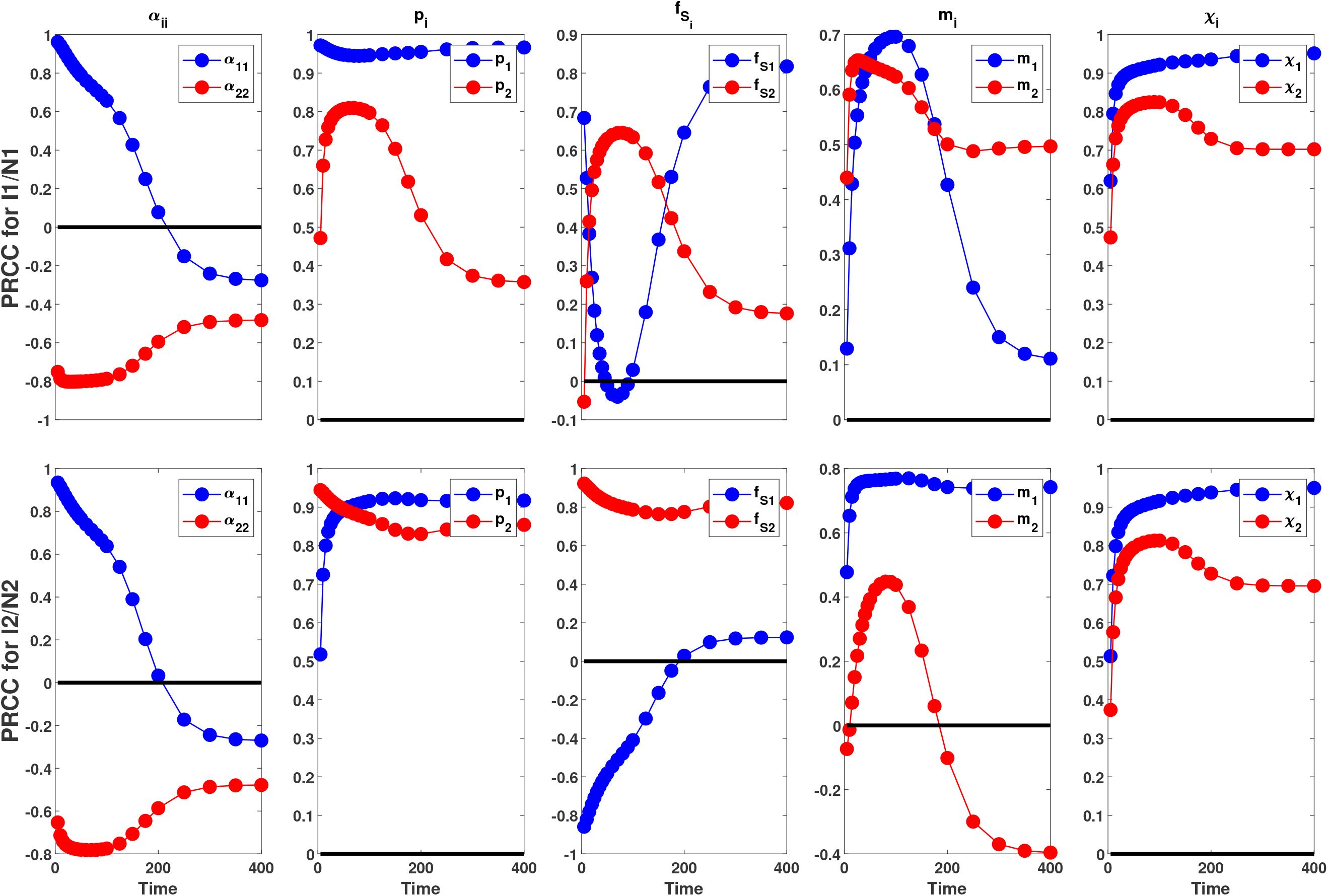
Comparison of the global sensitivities to all ecological parameters in Case 1. Row 1 shows the sensitivities for disease prevalence in host 1 (*I*_1_/*N*_1_) and row 2 shows the sensitivities for disease prevalence in host 2 (*I*_2_/*N*_2_). In each panel, solid blue circles denote the sensitivity to a parameter for host 1 (e.g., *α*_11_ or *p*_1_) and the solid red circles denote the sensitivity to a parameter for host 2 (e.g., *α*_22_ or *p*_2_). The solid black line denotes a PRCC value of 0. When comparing panels in the same column, the magnitudes and temporal patterns of the sensitivities are predicted by (i) intraspecific vs. interspecific processes when the solid blue curve in the top panel qualitatively matches the solid red curve in the bottom panel and the solid red curve in the top panel qualitatively matches the solid blue curve in the bottom panel or (ii) species’ identities when the solid blue curves in the top and bottom panels qualitatively match and the solid red curves in the top and bottom panels qualitatively match. See section 5.3.1 for additional details.

**Figure 7:**
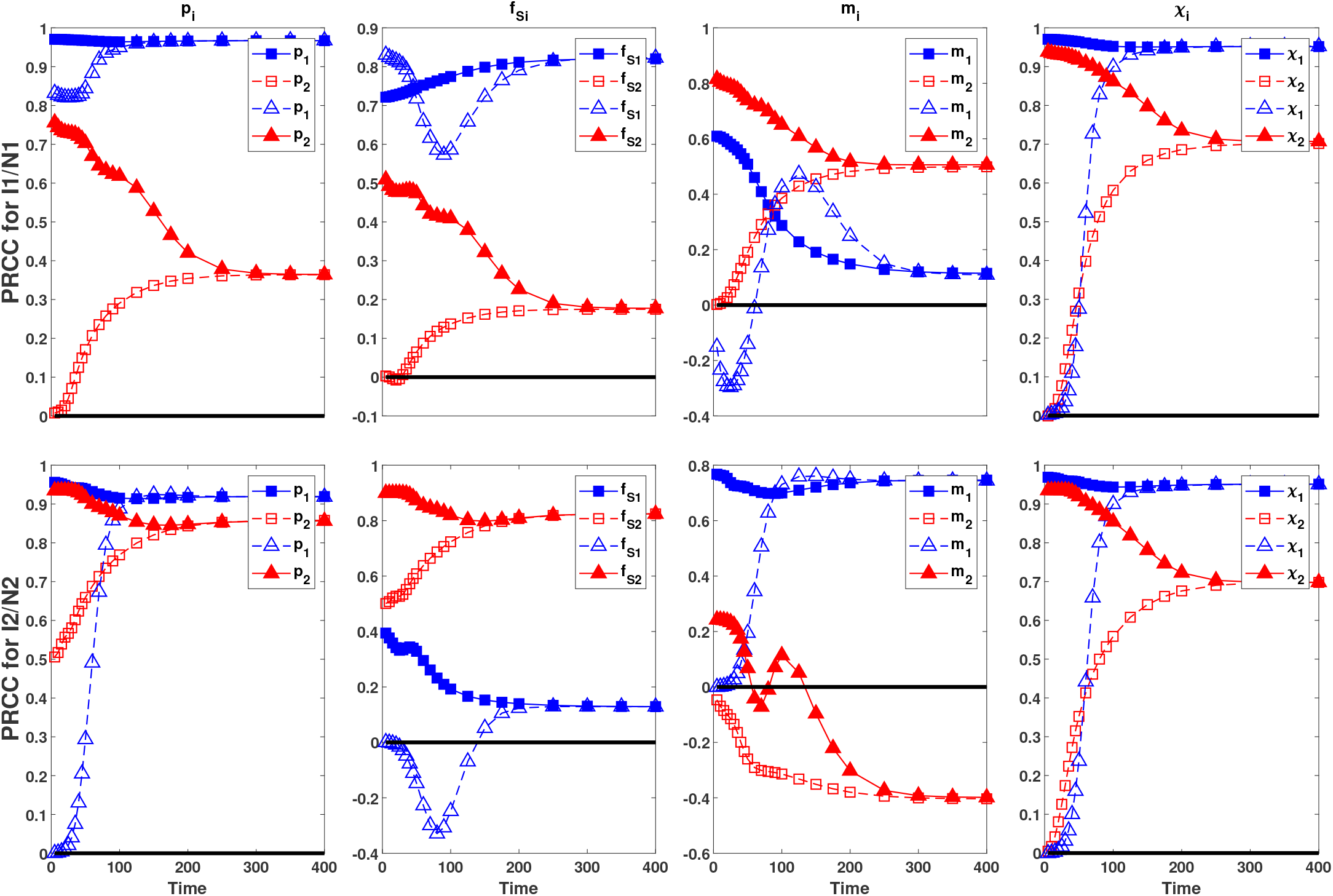
Comparison of the global sensitivities to all epidemiological parameters in Cases 2 and 3. Row 1 shows the sensitivities for disease prevalence in host 1 (*I*_1_/*N*_1_) and row 2 shows the sensitivities for disease prevalence in host 2 *I*_2_/*N*_2_). In each panel, blue denotes parameters for host 1, red denotes parameters for host 2, squares denote sensitivities for Case 2, triangles denote sensitivities for Case 3, solid lines with filled symbols denote parameters for the resident host (host 1 in case 2 and host 2 in case 3), and dashed lines with open symbols denote parameters for the invading host (host 2 in case 2 and host 1 in case 3). The solid black line denotes a PRCC value of 0. When comparing panels in the same column, the magnitudes and temporal patterns of the sensitivities are predicted by (i) intraspecific vs. interspecific processes when the solid blue curve with a filled shape in the top panel qualitatively matches the dashed red curve with the same open shape in the bottom panel, and the dashed blue curve with an open shape in the top panel qualitatively matches the solid red curve with the same filled shape in the bottom panel; (ii) the species’ roles of invader vs. resident when the solid blue curve with a filled shape in the top panel qualitatively matches the solid red curve with the same filled shape in the bottom panel, and the dashed blue curve with an open shape in the top panel qualitatively matches the dashed red curve with the same open shape in the bottom panel; and (iii) species’ identities when solid curves with filled symbols of one color in the top panel qualitatively match solid curves with filled symbols of the same color in the bottom panel, or when dashed curves with open symbols of one color in the top panel qualitatively match dashed curves with open symbols of the same color in the bottom panel. See section 5.3.2 for additional details.

**Figure 8:**
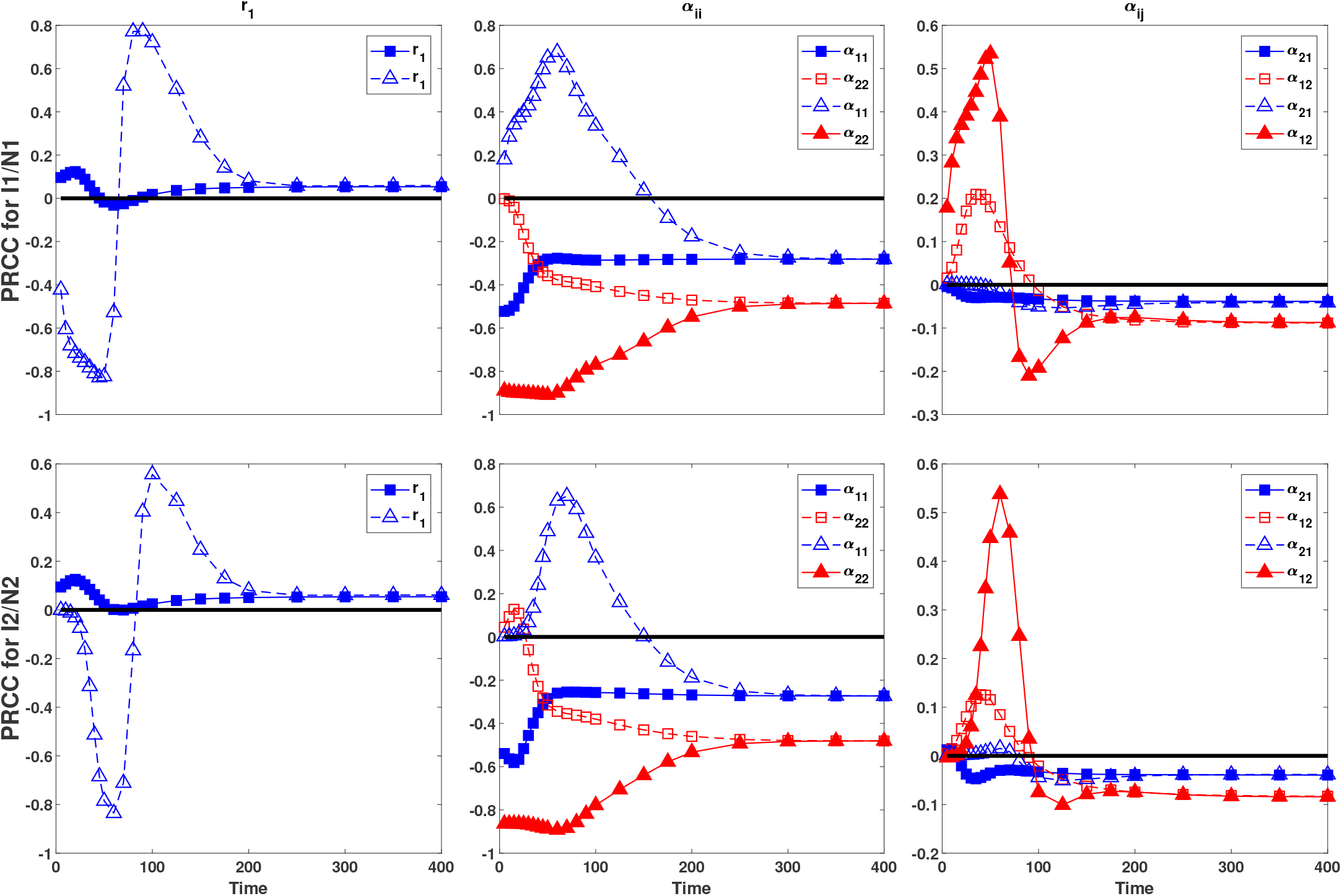
Comparison of the global sensitivities to all ecological parameters in Cases 2 and 3. Row 1 shows the sensitivities for disease prevalence in host 1 (*I*_1_/*N*_1_) and row 2 shows the sensitivities for disease prevalence in host 2 (*I*_2_/*N*_2_). In each panel, blue denotes parameters for host 1, red denotes parameters for host 2, squares denote sensitivities for Case 2, triangles denote sensitivities for Case 3, solid lines with filled symbols denote parameters for the resident host (host 1 in case 2 and host 2 in case 3), and dashed lines with open symbols denote parameters for the invading host (host 2 in case 2 and host 1 in case 3). The solid black line denotes a PRCC value of 0. When comparing panels in the same column, the magnitudes and temporal patterns of the sensitivities are predicted by (i) intraspecific vs. interspecific processes when the solid blue curve with a filled shape in the top panel qualitatively matches the dashed red curve with the same open shape in the bottom panel, and the dashed blue curve with an open shape in the top panel qualitatively matches the solid red curve with the same filled shape in the bottom panel; (ii) the species’ roles of invader vs. resident when the solid blue curve with a filled shape in the top panel qualitatively matches the solid red curve with the same filled shape in the bottom panel, and the dashed blue curve with an open shape in the top panel qualitatively matches the dashed red curve with the same open shape in the bottom panel; and (iii) species’ identities when solid curves with filled symbols of one color in the top panel qualitatively match solid curves with filled symbols of the same color in the bottom panel, or when dashed curves with open symbols of one color in the top panel qualitatively match dashed curves with open symbols of the same color in the bottom panel. See section 5.3.2 for additional details.

First, if the magnitudes and dynamics of the sensitivities are predicted by intraspecific versus interspecific processes, then either the sensitivities of *I_i_*/*N_i_* (*i* = 1, 2) to parameter *γ_i_* will be qualitatively similar or the sensitivities of *I_i_*/*N_i_* (*i* = 1, 2) to parameter *γ_j_* (*j* ≠ *i*) will be qualitatively similar. In such cases, the magnitudes or dynamics of the sensitivities can be predicted solely by knowing if the parameter corresponds to a process for the host of interest (i.e., an intraspecific process) or the other host species (i.e., an interspecific process). With respect to the figures, this means the patterns can be predicted by whether the focal parameter has the same color or the opposite color as the host of interest (where host 1 is blue and host 2 is red). For example, early in time in Case 1, prevalence in host *i* is more sensitive to its own probability of infection (*p_i_*; intraspecific parameter) than the probability of infection of the other host (*p_j_*, *j* ≠ *i*; interspecific parameter) (solid blue curve above solid red curve in top panel of column 2 in Figure 6 and solid red curve above solid blue curve in bottom panel of column 2 in Figure 6 for small *t*). As another example, also early in time in Case 1, prevalence in host *i* is more sensitive to the other host’s mortality rate (*m_j_*, *i ≠ j*; interspecific parameter) than its own mortality rate (*m_i_*; intraspecific parameter) (solid red curve above solid blue curve in top panel of column 4 in Figure 6 and solid blue curve above solid red curve in bottom panel of column 4 in Figure 6 for small *t*).

Second, if the magnitudes and dynamics of the sensitivities are predicted by the species’ roles of invader versus resident, then the sensitivities of *I*_1_/*N*_1_ and *I*_2_/*N*_2_ to the resident host’s parameters will be qualitatively similar and the sensitivities to the invader host’s parameters will be qualitatively similar. In such cases, the relative magnitudes or dynamics of the sensitivities can be predicted solely by knowing which parameter corresponds to a process for the resident host versus the invading host. With respect to the figures, this means the patterns can be predicted by which parameter has a solid curve with filled symbols (the resident’s parameter) or a dashed curve with open symbols (the invader’s parameter). For example, early in time in Cases 2 and 3, *I*_1_/*N*_1_ and *I*_2_/*N*_2_ are both more sensitive to the resident’s intraspecific competition coefficient (*α_ii_*) than the invader’s competition coefficient. Specifically, for small *t* in both panels of column 2 of Figure 8, the solid blue curve with filled squares is larger in magnitude than the dashed red curve with open squares (i.e., Case 2) and the solid red curve with filled triangles is larger in magnitude than the dashed blue curve with open triangles (i.e., Case 3).

Third, if the magnitudes and dynamics of the sensitivities are predicted by the species’ identities, then the sensitivities of *I*_1_/*N*_1_ and *I*_2_/*N*_2_ to parameter *q*_1_ will be qualitatively similar and the sensitivities to *q*_2_ will be qualitatively similar. In such cases, the relative magnitudes or dynamics of the sensitivities can be predicted only by knowing which parameter corresponds to which species. With respect to the figures, this means the patterns can be predicted solely by the color of the curve for the parameter. For example, in Case 1, prevalence in both species is more sensitive to the burst size of host 1 (*χ*_1_) than host 2 (*χ*_2_) at all points in time (solid blue curves above solid red curves for all time points in both panels in the fifth column of Figure 6). Thus, the relative magnitudes of the sensitivities can be predicted only by knowing which burst size corresponds to which host species.

Dividing the parameters into these two categories yields the following biological insight. If the magnitudes and temporal patterns can be predicted by intraspecific versus interspecific processes or the species’ roles of invader versus resident, then those parameters correspond to biological processes whose effects on disease prevalence can be predicted independent of specific characteristics (e.g., competitive ability or competence) of the two host species. This would suggest that those biological processes have the same effect in all systems, regardless of which host species are present. In comparison, if the dynamics are predicted by the species’ identities, then those parameters correspond to processes whose effects on disease prevalence can be predicted only when the specific characteristics of the host species are known. This would suggest that the biological processes can have different effects in systems where different host species are present. Whether the magnitudes and dynamics of the sensitivities can be predicted by intraspecific versus interspecific processes, the species’ roles or the species’ identities may change over time. This information is also useful because it yields insight about how the effects of processes on disease prevalence vary as conditions within the system change.

#### 5.3.1 Analysis of Case 1: Do temporal patterns differ between host species when the pathogen is introduced into the community?

Here we analyze the temporal dynamics of the PRCC sensitivity indices when the pathogen is introduced into a disease-free two-host community. We compare the temporal dynamics of the sensitivities for host species 1 (top row in Figure 6) and host species 2 (bottom row in Figure 6). Panels in each column show the sensitivities to parameters for host species 1 (solid blue curves with filled circles) and host species 2 (solid red curves with filled circles). For each parameter, the magnitudes and dynamics of the sensitivities are predicted by intraspecific versus interspecific processes if either the sensitivities of *I_i_*/*N_i_* (*i* = 1, 2) to parameter *q_i_* are qualitatively similar or the sensitivities of *I_i_*/*N_i_* (*i* = 1, 2) to parameter *q_j_* (*j ≠ i*) are qualitatively similar. In Figure 6, this means that the patterns are predicted by whether the focal parameter has the same color or the opposite color as the quantity of interest (where host 1 is blue and host 2 is red). In comparison, the magnitudes and dynamics of the sensitivities are predicted by species’ identities if the sensitivities of *I*_1_/*N*_1_ and *I*_2_/*N*_2_ to parameter *q_i_* are qualitatively similar and the sensitivities to *q_j_* (*i* ≠ *j*) are qualitatively similar. In Figure 6, this means that the patterns for both quantities of interest are predicted solely by the color of the curve for the parameter.

##### Sensitivities to epidemiological parameters

First consider the temporal dynamics of the PRCC values for the most sensitive epidemiological parameters (*p_i_*, 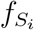, *χ_i_*, *m_i_*; columns 2-5 in Figure 6). Early in time (left side of each panel), the magnitudes of the sensitivities are predicted by intraspecific versus interspecific processes for *p_i_*, 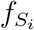, and *m_i_* and by the species’ identities for *χ_i_*. Specifically, prevalence in each species is more sensitive to the intraspecific probability of infection (*p_i_*; column 2), the intraspecific filtering rate (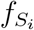; column 3), and the interspecific mortality rate (*m_j_*, *j* ≠ *i*; column 4), but prevalence in both species is more sensitive to the burst size of host 1 (*χ*_1_; column 5). Over time, the sensitivities transition to being predicted by intraspecific versus interspecific processes for 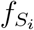 and *m_i_* and being predicted by species’ identities for *χ_i_* and *p_i_*. Specifically, at the end of the simulations (right side of each panel), each host is more sensitive to its own filtering rate 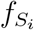 ; column 3) and the mortality rate of the other host (*m_j_*; column 4), and both hosts are more sensitive to the probability of infection for host 1 (*p*_1_; column 2) and the burst size of host 1 (*χ*_1_; column 5). Many of these transitions are non-monotonic, with each parameter having some period of time when the relative magnitudes are determined by the species’ identify (e.g., both hosts are more sensitive to the filtering rate of host 2 near *t* = 200; red curves are larger in magnitude than blue curves in column 3 of Figure 6). In addition, prevalence is both species is always more sensitive to *χ*_1_ than *χ*_2_ (blue curves above red curves in column 5).

The above yields three pieces of biological insight about systems where a pathogen is introduced into a disease-free community. First, during the initial stages of an outbreak, intraspecific versus interspecific processes determine how sensitive disease dynamics in each species is to processes directly affecting the increase (filtering, probability of infection) and decrease (mortality) of infected individuals in each host species. Second, however, for every process there exists periods of time later in the outbreak when the specific characteristics of the host species determine the relatively sensitivities. Third, the species’ identity and characteristics were always important for determining the relative sensitivity of the disease dynamics to burst sizes. Thus, while species’ identify and characteristics are less important during the initial stages of the outbreak, the species’ identify was always important for spore release and was important during some period of time for all other epidemiological processes.

##### Sensitivities to ecological parameters

Now consider the temporal dynamics of the PRCC values for the most sensitive ecological parameters (*α_ii_*; column 1 of Figure 6). The dynamics of the sensitivities are predicted by host identity. In particular, the sensitivities to the competition parameter for host 1 (*α*_11_) monotonically decrease from large positive values to small negative values (blue curves in column 1 of Figure 6). In comparison, the sensitivities to the competition parameter for host 2 (*α*_22_) are negative for all time, but transiently increase in magnitude before ultimately decreasing in magnitude (red curves in column 1 of Figure 6).

These patterns yield two pieces of biological insight about how species’ identities influences disease outbreaks. First, reducing the growth rate of host 1 and increasing the growth rate of host 2 (via reduced competition) facilitates faster increases in infections in both populations. This is due to host 2 having higher competence than host 1, where the competence of host *i* is measured by the ratio *χ_i_p_i_*/*m_i_δ* (Cortez, 2021). Specifically, faster growth is host 2 yields a faster increase in spore density, which leads to more infections in both host species. Second, the disease dynamics are more sensitive to competition at early and intermediate stages of the outbreak and less sensitive later in time. However, the specific points in time when sensitivity to competition in each species is maximal depends on the species identity (earlier in time for host 1 than host 2). Overall, this suggests that knowing each species’ characteristics is crucial for determining how intraspecific and interspecific host competition influence outbreak patterns.

#### 5.3.2 Analysis of Cases 2 and 3: How do temporal patterns depend on which host species is the resident versus invader?

Here we focus on similarities and differences in the temporal dynamics of the PRCC values when one host species is the resident and the other host species is the invader. We focus on (i) comparing the temporal dynamics of the sensitivities for the invading species (solid curves with filled symbols in Figures 7 and 8) and (ii) comparing the temporal dynamics of the sensitivities for the resident species (dashed curves with open symbols in Figures 7 and 8). Panels in each column show the sensitivities to parameters for host species 1 (blue curves) and host species 2 (red curves) in Case 2 (squares) and Case 3 (triangles). For each parameter, the magnitudes and dynamics of the sensitivities are predicted by intraspecific versus interspecific processes if the sensitivities of *I_i_*/*N_i_* (*i* = 1, 2) to parameter *q_i_* are qualitatively similar and the sensitivities of *I_i_*/*N_i_* (*i* = 1, 2) to parameter *q_j_* (*j* ≠ *i*) are qualitatively similar. In Figures 7 and 8, this means that the patterns are predicted by whether the focal parameter has the same color or the opposite color as the quantity of interest (where host 1 is blue and host 2 is red). The magnitudes and dynamics of the sensitivities are predicted by the species’ roles of invader versus resident if the sensitivities of *I*_1_/*N*_1_ and *I*_2_/*N*_2_ to the resident host’s parameters are qualitatively similar and the sensitivities to the invader host’s parameters are qualitatively similar. In Figures 7 and 8, this means that the patterns are predicted by whether the focal parameter has a solid curve with filled symbols (the resident’s parameters) or a dashed curve with open symbols (the invader’s parameters). Finally, the magnitudes and dynamics of the sensitivities are predicted by species’ identities if the sensitivities of *I*_1_/*N*_1_ and *I*_2_/*N*_2_ to parameter *q_i_* are qualitatively similar and the sensitivities to *q_j_* (*i* ≠ *j*) are qualitatively similar. In Figures 7 and 8, this means that the patterns for both quantities of interest are predicted solely by the color of the curve for the parameter.

##### Sensitivities to epidemiological parameters

First consider the temporal dynamics of the PRCC values for the most sensitive epidemiological parameters (*p_i_*, 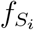, *χ_i_*, *m_i_*); see Figure 7. Early in time in both cases 2 and 3, the species’ roles (i.e., invader versus resident) determine if prevalence in each species is more sensitive to the resident’s or invader’s parameters. Specifically, both hosts are more sensitive to the resident’s epidemiological parameters than the invader’s parameters (solid curve larger in magnitude than dashed curves early in time in Figure 7); the only exception is that the invading host is more sensitive to its filtering rate 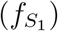 than the filtering rate of the resident host species (blue curves above red curves in top panel of column 2 in Figure 7 and red curves above blue curves in bottom panel of column 2). Over time, the sensitivities transition to being predicted by intraspecific versus interspecific processes for 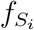 and *m_i_* and being predicted by species’ identities for *χ_i_* and *p_i_*. Many of these transitions are monotonic (e.g., columns 1 and 4 in Figure 7), however non-monotonic changes occur for the filtering rates (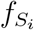; column 2 of Figure 7) and mortality rates (*m_i_*; column 3 in Figure 7).

Two pieces of biological insight are gained from the above. First, early in time, the magnitudes of the sensitivities align with what one would expect based on the host densities. Specifically, when either host species is introduced into the community, the initial disease dynamics are more sensitive to the epidemiological traits of the resident because it is at higher abundance than the invader. Thus, early in time, the species’ roles determine the relative sensitivity to epidemiological processes in each host. Second, later in time, the sensitivities of the disease dynamics depend on the specific characteristics (i.e, parameter values) of each host species, and the magnitudes of the sensitivities cannot necessarily be explained by the host densities. For example, the larger magnitudes for *p*_1_ than *p*_2_ and for *χ*_1_ than *χ*_2_ align with host 1 being at higher density than host 2, but the temporal and long-term dynamics for 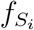 and *m_i_* do not align. Overall, this shows that the epidemiological dynamics of the system are not strongly sensitive to host competence and identity early in time after a host introduction, but can be strongly sensitive later in time after an introduction.

##### Sensitivities to ecological parameters

Now consider the temporal dynamics of the PRCC values for the most sensitive ecological parameters (*α_ij_*, *r_i_*); see Figure 8. Early in time in both cases 2 and 3, the species’ roles (i.e., invader versus resident) determine if prevalence in each species is more sensitive to the resident’s or invader’s parameters. Specifically, both hosts are more sensitive to the resident’s competition parameters than the invader’s (solid curves with filled symbols are larger in magnitude than dashed curves with the same open symbol and the opposite color early in time in Figure 8). Later in time, the magnitudes of the sensitivities depend on the species’ identities, specifically which host is the invader and resident. When host 1 is the resident (Case 2), the sensitivities for all competition parameters are below the 0.6 threshold for all time (all curves with squares in Figure 8 are less than 0.6 in magnitude for all time). In comparison, when host 2 is the resident (Case 3), the sensitivities to the exponential growth rate of host 1 (*r*_1_) and both intraspecific competition parameters (*α*_11_, *α*_22_) are above the threshold for some period of time (curves with triangles in columns 1 and 2 of Figure 8 are greater than 0.6 in magnitude). In addition, the sensitivities to the interspecific competition parameters (*α*_12_, *α*_21_) are often larger in Case 3 than Case 2 (curves with triangles are typically greater in magnitude than curves with squares in column 3 of Figure 8).

Two pieces of biological insight are gained from the above. First, early in time, the magnitudes of the sensitivities align with what one would expect based on the host densities. Specifically, when either host species is introduced into the community, the initial disease dynamics are more sensitive to resident’s parameters because it is at higher abundance than the invader. Thus, early in time, the species’ roles determine the relative sensitivity to ecological processes affecting each host. Second, whether the epidemiological dynamics are sufficiently sensitive to competition (defined as magnitudes greater than 0.6) depends on the specific identities of the invader and resident. Specifically, the prevalence dynamics are much more sensitive when host 1 is the invader (curves with triangles) than when host 2 is the invader (curves with squares). Thus, the specific identities of the resident and invader determine if intraspecific and interspecific host competition have strong effects on disease dynamics. Interestingly, this pattern is the opposite of what one might expect given that host 2 is the stronger intraspecific and interspecific competitor (*α_i_*_2_ larger than *α_i_*_1_). We suspect that it is due to host 2 being at lower densities than host 1, which results in competition having a relatively larger effect on the disease dynamics of the system.

## 6 Discussion and Conclusions

In this study we used global sensitivity analysis to gain insight into how ecological and epidemiological processes shape the temporal dynamics of a two-host-one-pathogen community. We focused on three biologically important scenarios, defined by the introduction of the pathogen into a disease-free community or the introduction of a second host species into an endemic single-host community. Our numerical results show that the signs and magnitudes of some effects can be predicted by factors independent of the species’ identities (intraspecific versus interspecific processes; invader versus resident) whereas others can only be predicted by knowing the species’ identities and specific characteristics (i.e., their competitive abilities and disease competence). Moreover, whether the effects are predicted by the species’ roles versus the species’ identities can change over time. Below we discuss the biological insights gained from this approach and how our results add to the growing body of theory on how host biodiversity shapes disease dynamics.

Global sensitivity analysis is a well developed methodology that has been widely applied in engineering settings where understanding reliability is very important (Saltelli, 2002; Saltelli et al., 2005; Wu, 1994). Within the past few decades, sensitivity analysis has begun to be used to assess model predictions, experimental design, and robustness in a variety of biological applications (Blower and Dowlatabadi, 1994; Jarrett et al., 2015, 2017b; Renardy et al., 2019; Aggarwal et al., 2019, 2021; Marino et al., 2008; Wentworth et al., 2016; Hanthanan Arachchilage and Hussaini, 2021). One of the differences between more traditional applications and biological applications is in the interpretation of the rankings. Often, the goal of engineering applications is to use GSA to remove parameters that are insensitive while biological insight is typically driven by which parameters are sensitive. Highlighting parameters into those that dominate outcomes helps develop insight into experimental design, interpretation of the model, and robustness of conclusions. Additionally, since biological applications are typically dynamic, analysis of the time evolution of GSA rankings also helps understand which processes matter most, when and with respect to what observations.

This study demonstrates that the temporal patterns of the sensitivities can yield insight into how specific processes are affecting quantities of interest at different points in time. In particular, comparing the temporal dynamics of the sensitivities of the two host species identifies when the effects of a species process are driven by the specific characteristics of each species (i.e., species’ identities) versus properties of the system that are independent of the species’ identities (i.e., intraspecific versus interspecific processes and the species’ roles of invader versus resident). Biologically, this is useful because it helps determine what information is needed to make predictions about how a specific process shapes disease dynamics. For example, when a pathogen is introduced into a disease-free community (Case 1), one must know the competitive abilities of each host species (e.g., *α_ij_*) to predict how host competition affects disease dynamics at all points time. In contrast, the signs and magnitudes of the effects of infection, spore release, and mortality on disease dynamics early in time are predicted only by knowing whether the process is affecting conspecifics (i.e., intraspecific processes) or heterospecifics (i.e., interspecific processes). While these particular comparisons are possible because of the symmetry of the model (the system is structurally invariant to changing the labels of host species 1 and 2), analogous comparisons in other ecological and epidemiological models may yield additional insight about how specific processes shape system dynamics.

Many studies have argued that the relationships species richness (i.e., the number of host species) and disease levels in a focal host are likely to be context dependent (LoGiudice et al., 2008; Randolph and Dobson, 2012; Rohr et al., 2020; Halliday et al., 2020). Previous empirical studies (Telfer et al., 2005; Dizney and Ruedas, 2009; Searle et al., 2016; Levine et al., 2017; Hydeman et al., 2017; Zimmermann et al., 2017; Luis et al., 2018) and theoretical studies (Dobson, 2004; Rudolf and Antonovics, 2005; Joseph et al., 2013; Mihaljevic et al., 2014; O’Regan et al., 2015; Searle et al., 2016; Faust et al., 2017; Roberts and Heesterbeek, 2018; Cortez and Duffy, 2021; Cortez, 2021) have shown that whether the addition of a host species leads to increased or decreased disease in a focal host depends on the specific characteristics (e.g., competitive ability or disease competence) of the host species that are present in the community and added to the community. The time varying signs and magnitudes of the global sensitivities in this study suggest that the context dependent relationships may also depend on the time scale of interest. However, we note that our results are not direct evidence of time varying host richness-disease relationships because we do not look at the relationship between disease prevalence in host *i* and the density of host *j* (*j* ≠ *i*), i.e., the sensitivity of prevalence in host *i* to host *j* density. Nonetheless, the sensitivities computed in this study are related and prior studies (Cortez and Duffy, 2021; Cortez, 2021) have shown that local sensitivities to parameters can yield some insight about local sensitivities to host densities.

Related to the issue of time varying richness-disease relationships, prior empirical studies (Evans and Entwistle, 1987; Johnson et al., 2008; Searle et al., 2011; Orlofske et al., 2012; Becker et al., 2014; Venesky et al., 2014; Hopkins et al., 2020) have used short term exposure experiments to infer how the presence or absence of alternative host species affect disease levels in a focal host species. The intuition is that higher (lower) prevalence in a short term experiment will translate to higher (lower) prevalence over the long term. The sensitivities for the host filtering rates 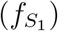 in Case 1 suggest that this intuition may be incorrect. Specifically, introduction of a host with a higher filtering rate could lead to lower infection prevalence over the short term (because the sensitivity of *I_i_*/*N_i_* to 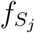 for *i* ≠ *j* is negative for small *t*) but positive over the long term (because the sensitivity of *I_i_*/*N_i_* to 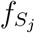 for *i* ≠ *j* is positive for small *t*); see Table 2. The difference arises because the short-term dynamics do not account for indirect effects between species that become larger in magnitude over time. In particular, increased uptake leads to more infections in host *j*, which leads to greater release of spores by infected individuals of host *j* and ultimately increased infection rates in host *i*. Our results also show that for some processes the effects on disease prevalence have effects of constant sign. For example, increased burst sizes for either host (larger *χ_i_*) has a positive effect on disease prevalence in both host species at all time points. Thus, while our results suggest caution when making inferences about long term disease levels from short-term experiments, they also illustrate that global sensitivity analysis can be used to make predictions about when processes may potentially have effects of different signs on system dynamics.

In addition to making predictions about when processes may potentially have effects of different signs at different time points, sensitivity analysis can also allow one to make predictions about whether those differences in sign are likely to alter the dynamics of the system. In particular, if a quantity of interest is weakly sensitive to a specific parameter or process, then sign changes in the sensitivity for that parameter is less likely to have a measurable effect on the system dynamics. More generally, ranking the sensitivities of all parameters allows one to identify the most sensitive parameters and reduce the number of parameters that need to be considered. Returning to the filtering rate and burst size examples from above, the sensitivity of prevalence in host 1 (*N*_1_/*I*_1_) to host 2’s filtering rate 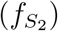 is below our threshold for all points in time whereas the sensitivity of prevalence in host 2 (*I*_2_/*N*_2_) to host 1’s filtering rate 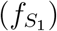 is above the threshold for intermediate time points. This suggests that short-term predictions about the effects of host 2 on disease prevalence in host 1 are less likely to qualitatively differ from the long-term predictions. In comparison, short-term predictions about the effects of host 1 on disease prevalence in host 2 are more likely to qualitatively differ from the long-term predictions, because prevalence in host 2 is more sensitive to the filtering rate of host 1.

While this study shows that global sensitivity analysis can be a useful tool, more can be done to gain further insight. First, we have applied our approach to a single region of parameter space, and it possible that different patterns would be observed in other regions. Indeed, previous studies (Roberts and Heesterbeek, 2018; Cortez and Duffy, 2021; Cortez, 2021) have shown that the signs of local sensitivities at equilibrium can differ across parameter space. Exploring how the temporal patterns of the sensitivities vary across parameter space is likely to yield insight about if and when patterns can be generally explained by the species’ identities or by other factors (e.g., invader versus resident or intraspecific versus interspecific processes).

Second, additional insight could also be gained by applying this approach to other metrics of disease. Common metrics include the basic reproduction number (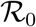; Dobson 2004; Joseph et al. 2013; Mihaljevic et al. 2014; O’Regan et al. 2015), the density of infected individuals (*I_i_* in our model; Roche et al. 2012; Searle et al. 2016) and disease prevalence (*I_i_*/*N_i_* in our model; Rudolf and Antonovics 2005; Roche et al. 2012; Strauss et al. 2015; Faust et al. 2017; Cortez and Duffy 2021; Cortez 2021). Prior work has shown that predictions about host richness-disease relationships differ between metrics (e.g., introduction of a host can lead to higher 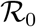 and lower prevalence at equilibrium) (Roche et al., 2012; Roberts and Heesterbeek, 2018; Cortez and Duffy, 2021). Applying global sensitivity analysis to these other metrics could yield additional insight into the different effects ecological and epidemiological processes have on disease dynamics over different time scales. For example, our partitioning of effects could also be useful when exploring how the presence of other species in a community (e.g., predators) affect disease dynamics in a focal host species.

Third, additional insight could be gained by using alternative global sensitivity measures. One alternative is Sobol’ sensitivity indices, where the first order Sobol’ index (*S_i_*) estimates the reduction in variance in the quantity of interest that occurs when parameter *q_i_* is held fixed. By definition, Σ_*i*_ *S_i_* ≤ 1. A system is said to be an additive system when Σ_*i*_ *S_i_* = 1. When Σ_*i*_ *S_i_* ≪ 1, variation in the quantity of interest is due to higher order parameter interactions. We estimated the Sobol’ first order indices for the quantities of interest and region of parameter space analyzed in this study and found that Σ_*i*_ *S_i_* ≈ 1. This suggests that higher order interactions between parameters only have small effects on the quantities of interest. Because PRCC and Sobol’ first order indices provide similar parameter rankings, our results from the Sobol’ sensitivity indices suggest that the PRCC indices are sufficient to explore the dynamics of the system completely. However, in systems where Σ_*i*_ *S_i_* ≪ 1 (e.g., the epidemiological system in Hanthanan Arachchilage and Hussaini (2021)), higher order sensitivity indices can provide new insight into how interactions between processes influence system behavior.

Fourth, PRCC and Sobol’ sensitivity indices assume that the input parameters, *q_i_*, are statistically uncorrelated. However, correlations between parameters can arise in many systems. For example, in this system susceptible individual filtering rates, infected individual filtering rates, and the competition coefficients are all likely to be positively correlated. Variance based approaches, such as those in (Kucherenko et al., 2012) and (Mara and Tarantola, 2012), can be used to account for correlated parameters. Accounting for correlations may yield additional insight about the sensitivities to specific processes and the relationships between first and higher order indices.

In closing, this study illustrates how GSA methodology can provide useful insight into the ecological and epidemiological processes that are modeled using mathematics. Understanding the details of specific quantities of interest, time-evolution, and interpretation of the GSA rankings requires understanding of the underlying mathematics, modeling, and biology, and the synthesis provides more information that each aspect alone.

## 7 Acknowledgements

MHC was supported by the National Science Foundation under Award DEB-2015280.

## Appendix

## A Local Sensitivities at equilibrium

The following sections briefly present the analytical method for computing the local sensitivities at equilibrium and then apply the theory in order to identify the different indirect pathways affecting each sensitivity and explain the signs of the associated indirect effects. The method and results are based off of the work in Cortez and Duffy (2021) and Cortez (2021), who analyzed generalized versions of models (1)-(5) and (6)-(8).

## A.1 Method for analytically computing local sensitivities at equilibrium

Consider a system of equations *dx_i_/dt* (1 ≤ *i* ≤ *n*) with an equilibrium 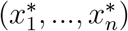 and parameter *γ*. The local sensitivity of 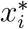 with respect to *γ* is

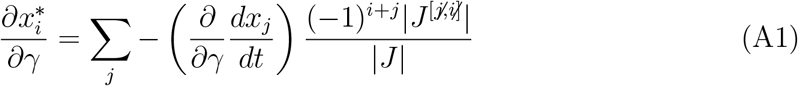

where *J* is the Jacobian of the system evaluated at the equilibrium, *J* ^[*j/, /i*]^ is the submatrix of *J* where row *j* and column *i* have been removed, and |*M*| denotes the determinant of matrix *M* .

For model (6)-(8), the Jacobian evaluated at the endemic equilibrium 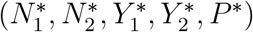 is

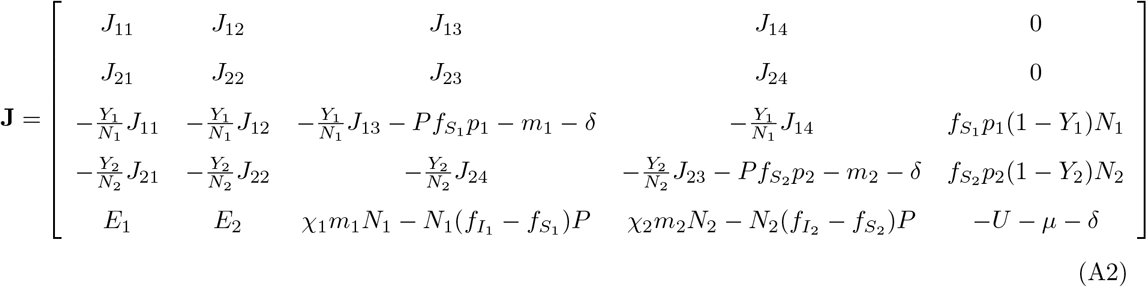

where the order of the variables is *N*_1_-*N*_2_-*Y*_1_-*Y*_2_-*P* and

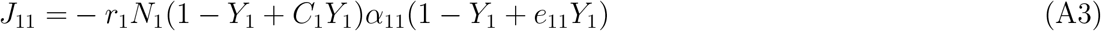

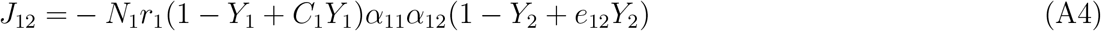

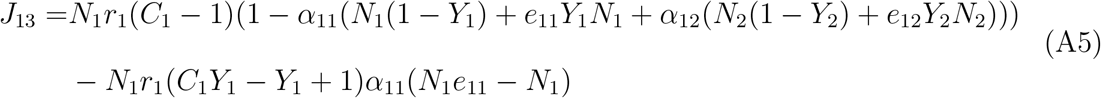

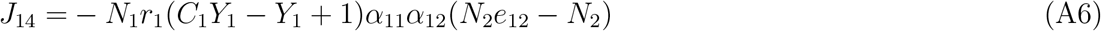

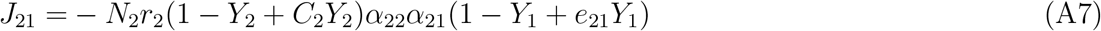

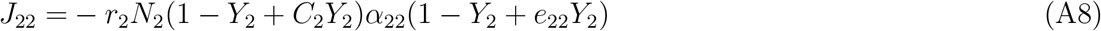

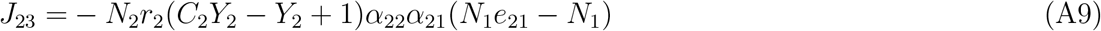

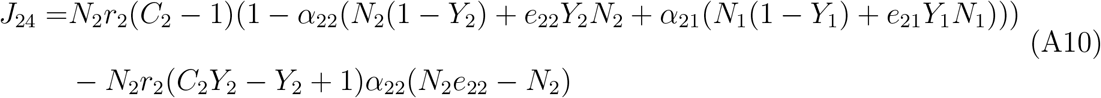

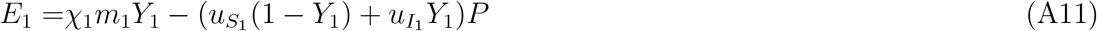

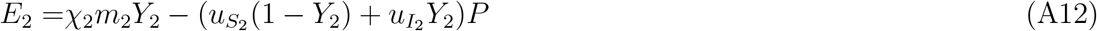

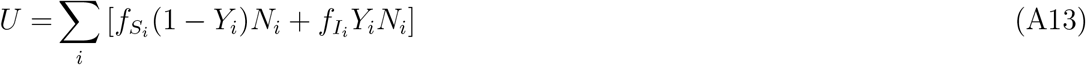

The way to biologically interpret equation (A1) is the following. Each Jacobian entry corresponds to sets of ecological and epidemiological processes. For example, the entry *J*_35_ = (*∂/∂P*)(*dY*_1_/*dt*) defines how the growth rate of infection prevalence in host population 1 (*dY*_1_/*dt*) increases as spore density (*P*) increases. 2. Each entry of the sum in equation (A1) defines how a change in parameter *q* affects the growth rate of variable *x_j_* and how those effects propagate through the community and ultimately affect the equilibrium density of variable *x_i_*. The propagating effects are indirect effects of parameter *γ* on 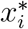. They can be visualized as an indirect pathway where arrows denote the direct effects of a parameter or variable on the dynamics of a variable, and the sign of each direct effect is computed using a partial derivative. For example, the local sensitivity of equilibrium disease prevalence in host 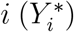 with respect to the burst size of host *j* (*χ_j_*) is defined by the derivative 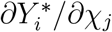. For this particular sensitivity, equation (A1) reduces to a single term representing the indirect effect of *χ_i_* on 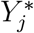. The corresponding indirect pathway is *χ_i_* → *P* → *Y_j_* where the signs of the direct effects *χ_i_* → *P* and *P* → *Y_j_* are positive because 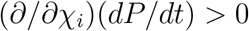 and (*∂/∂P*)(*dY_j_/dt*) > 0, respectively.

## A.2 Interpretations of local sensitivities at equilibrium

Evaluating the Jacobian (A2) at the endemic equilibrium for the nominal parameter values and applying the theory in Cortez (2021) yields the following predictions about the signs of the local sensitivities at equilibrium; see “LSA Eq.” columns in Table 2 for a summary.

## Host filtering rates 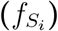 and probabilities of infection (*p_i_*)

The local sensitivity of infection prevalence in each host species to each parameter is positive. The signs of the sensitivities are determined by two kinds of indirect pathways: (i) *q_i_* → *Y_i_* and (ii) *q_i_* → *Y_i_* → *N_k_* → *P* → *Y_j_* (*j* ≠ *i*, *k* ∈ {1, 2}), where *q_i_* is a parameter in the *dY_i_*/*dt* equation. The first pathway accounts for how changes in the infection rate for host *i* affects disease prevalence in host *i* (*q_i_* → *Y_i_*). The indirect effect for the first pathway is always positive because increased infection rates lead to higher equilibrium prevalence. The second kind of pathway accounts for how changes in disease prevalence in host *i* affect intraspecific and interspecific competition (*Y_i_* → *N_k_*), changes in the density of host *k* affect spore release by host *k* (*N_k_* → *P*), and changes in spore density affect disease prevalence in the other host species (*P* → *Y_j_*). In these pathways, the direct effect *N_k_* → *P* is positive if host *k* is a net source of spores (defined by 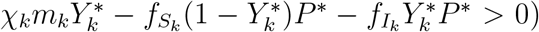 and negative if host *k* is a net sink of spores (defined by 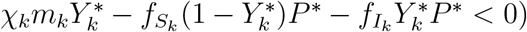. In general, the indirect effects for the second pathway can be positive or negative. For the nominal parameter set, the direct effects *N_k_* → *P* are positive, which results in all pathways having positive indirect effects. Consequently, the local sensitivities for the host infection rates (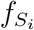, *p_i_*) are positive for both quantities of interest.

## Host burst sizes (*χ_i_*)

The local sensitivity of prevalence in each host species to each parameter is positive. The signs of these sensitivities are determined by the indirect pathways *χ_i_* → *P* → *Y_j_*. Because both direct effects are positive, the indirect effect and local sensitivities are always positive.

## Infected host morality rates (*m_i_*)

The local sensitivity of prevalence in each host species to the mortality rate of host 1 (*m*_1_) is positive. The local sensitivity of prevalence in host 1 to the mortality rate of host 2 (*m*_2_) is also positive, but the local sensitivity of prevalence in host 2 to the mortality rate of host 2 is negative. Instead of listing the many pathways that determine these sensitivities, we focus on the two most important pathways: (i) *μ_i_* → *Y_i_* and (ii) *μ_i_* → *P* → *Y_j_*. The first pathway accounts for how increased mortality of infected individuals decreases infection prevalence; the indirect effect for this pathway is negative. The second pathway accounts for how increased mortality leads to increased release of spores (*μ_i_* → *P*), which in turn affects disease prevalence in each host species (*P* → *Y_j_*); the indirect effect for this pathway is positive. The local sensitivities *∂Y_i_/∂m_j_* (*i ≠ j*) are positive because they are primarily determined by the second pathway. The local sensitivity *∂Y*_1_/*∂m*_1_ is positive because the second pathway is larger in magnitude than the first. In contrast, the local sensitivity *∂Y*_2_/*∂m*_2_ is negative because the first pathway is larger in magnitude than the second. The reason for the difference in sign is that host 2 has a much smaller equilibrium density than host 1. This causes increased mortality of host 2 (*m*_2_) to have a much larger direct effect on disease prevalence in the second host (*Y*_2_). In total, this results in positive local sensitivities for the host mortality rates, except for the local sensitivity of disease prevalence in the second host to its mortality rate.

## Reproduction rates (*r_i_, c_i_*) and competition parameters (*α_ij_*, *e_ij_*)

The local sensitivity of prevalence in each host species to each host reproduction rate parameter (*r_i_, c_i_*) is positive. The local sensitivity of prevalence in each host species to each competition parameter (*α_ij_*, *e_ij_*) is negative. The signs of the sensitivities are determined by the two indirect pathways (i) *q_i_* → *N_i_* → *P* → *Y_j_* and (ii) *q_i_* → *N_i_* → *N_k_* → *P* → *Y_j_* where *k* ≠ *i* and *q_i_* is a parameter in the *dN_i_/dt* and *dY_i_*/*dt* equations. The first pathway accounts for how changes in the parameter affect the density of host *i* (*q_i_* → *N_i_*), changes in density affect the production of spores by host *i* (*N_i_* → *P*), and change in spore density affect disease prevalence in both host species (*P* → *Y_j_*). The second pathway accounts for how changes in the density of host *i* affect the density of host *k* (*N_i_* → *N_k_*), changes in the density of host *k* affect spore density (*N_k_* → *P*), and changes in spore density affect prevalence in both host species (*P* → *Y_j_*). In general, the indirect effects for the two pathways can be positive or negative and have the same or different signs. For the nominal parameter values, the direct effects *N_i_* → *P* and *N_k_* → *P* are both positive, which results in the indirect effects for the two pathways having opposite signs. Because the indirect effect for the first pathway is larger in magnitude, the local sensitivities for the host reproduction rates (*r_i_, c_i_*) are positive and the local sensitivities for the competition parameters (*α_ij_*, *e_ij_*) are negative for both quantities of interest.

## Infected host uptake rates 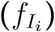 and the spore degradation rate (*μ*)

The local sensitivity of prevalence in each host species to each parameter is negative. The signs of the local sensitivities are determined by the indirect pathways *γ* → *P* → *Y_i_* where *γ* is one of the parameters. The indirect effects and local sensitivities are negative because the direct effect *γ* → *P* is negative.

## A.3 Comparison of signs of local sensitivities at equilibrium and derivatives of model equations

The signs of the derivatives of the model equations are summarized in the “Model Eqns.” columns of Table 2 and signs of the local sensitivities at equilibrium are summarized in the “LSA Eq.” columns in Table 2. The signs of the derivatives of the model equations agree with the signs of the local sensitivities for the infection probabilities (*p_i_*) and many of the parameters affecting the spore growth rate, including the filtering rates of infected individuals 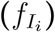, the spore burst sizes (*χ_i_*), the spore degradation rate (*μ*), and the destructive sampling rate (*δ*). The signs do not agree for the filtering rates of susceptible individuals 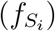, disease-induced mortality rates (*m_i_*) and all reproduction (*r_i_*, *c_i_*) and competition (*α_ij_*, *e_ij_*) parameters. The specific reasons for disagreement are the following.

First, the predictions for susceptible host filtering rates disagree because while increased filtering decreases the spore growth 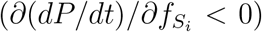, it also increases host infection rates 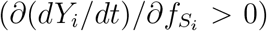, which can ultimately lead to increased spore densities due to greater rates of spore release. Second, the predictions for mortality rates differ because while increased mortality initially reduces disease prevalence (*Y_i_*) due to the removal of infected individuals (*∂*(*Y_i_/dt*)/*∂m_i_* < 0), it also increases rates of spore release, which can result in increased rates of transmission and disease prevalence. Third, the predicted signs for the reproduction and competition parameters differ because while increased reproduction reduces disease prevalence (*Y_i_*) due to an influx of susceptible individuals (e.g., *∂*(*Y_i_/dt*)/*∂r_i_* < 0), increased susceptible density results in increased rates of transmission and ultimately increased disease prevalence.

## B Detailed analysis of temporal changes in signs of global sensitivities

The following provides detailed explanations for why some parameters have global sensitivities with signs that change over time. To provide some extra insight, we discuss all parameters whose PRCC values exceed 0.6 for at least one quantity of interest at some point in time (i.e, max(|*PRCC*(*t*)|) ≥ 0.6). The rough trends are summarized in Table 2 of the main text. Time series for the global sensitivities for all parameters in each case are provided in each section.

## B.1 Case 1: Pathogen invades two-host disease-free equilibrium

In Case 1, the pathogen is introduced into a disease-free two-host system at equilibrium. Time series for the global sensitivities for all parameters are given in Figures B1 and B2. For many of the most influential parameters (*p_i_*, *χ_i_*, *m_i_*, *α*_2*j*_, *δ*, *μ*) the signs of the global sensitivities at all points in time agree with the signs of the local sensitivities; see appendix A.2 for interpretations. However, differences in signs occur for some parameters 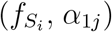 over short times scales because spore density is initially very low. Explanations for the differences are given below.

**Figure B1:**
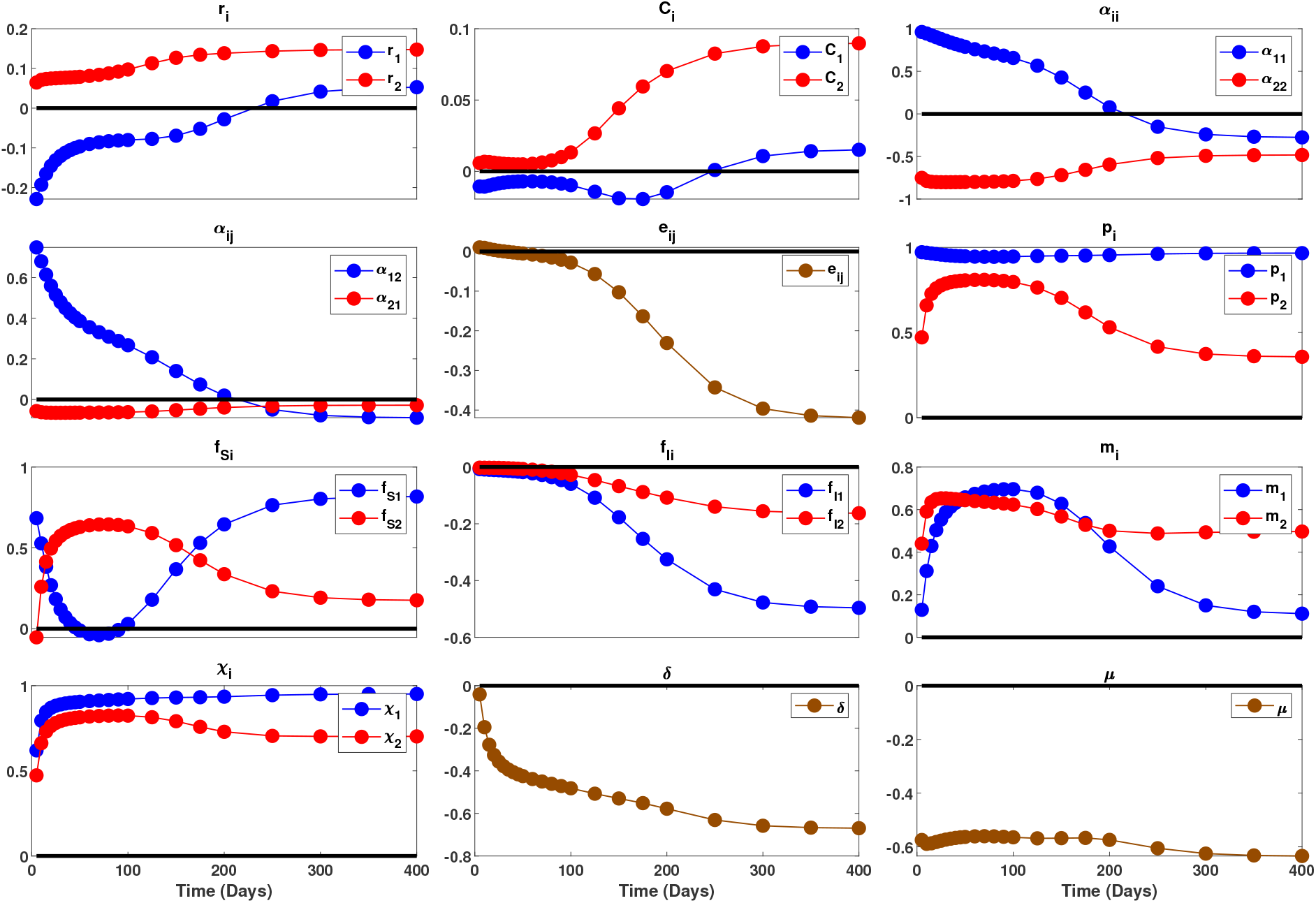
Global sensitivities of prevalence in host 1 (*I*_1_/*N*_1_) to all parameters in Case 1. In each panel, the blue curve corresponds to the PRCC values for a parameter for host 1, the red curve corresponds to a PRCC value for a parameter for host 2, and the black line denotes a value of zero.

**Figure B2:**
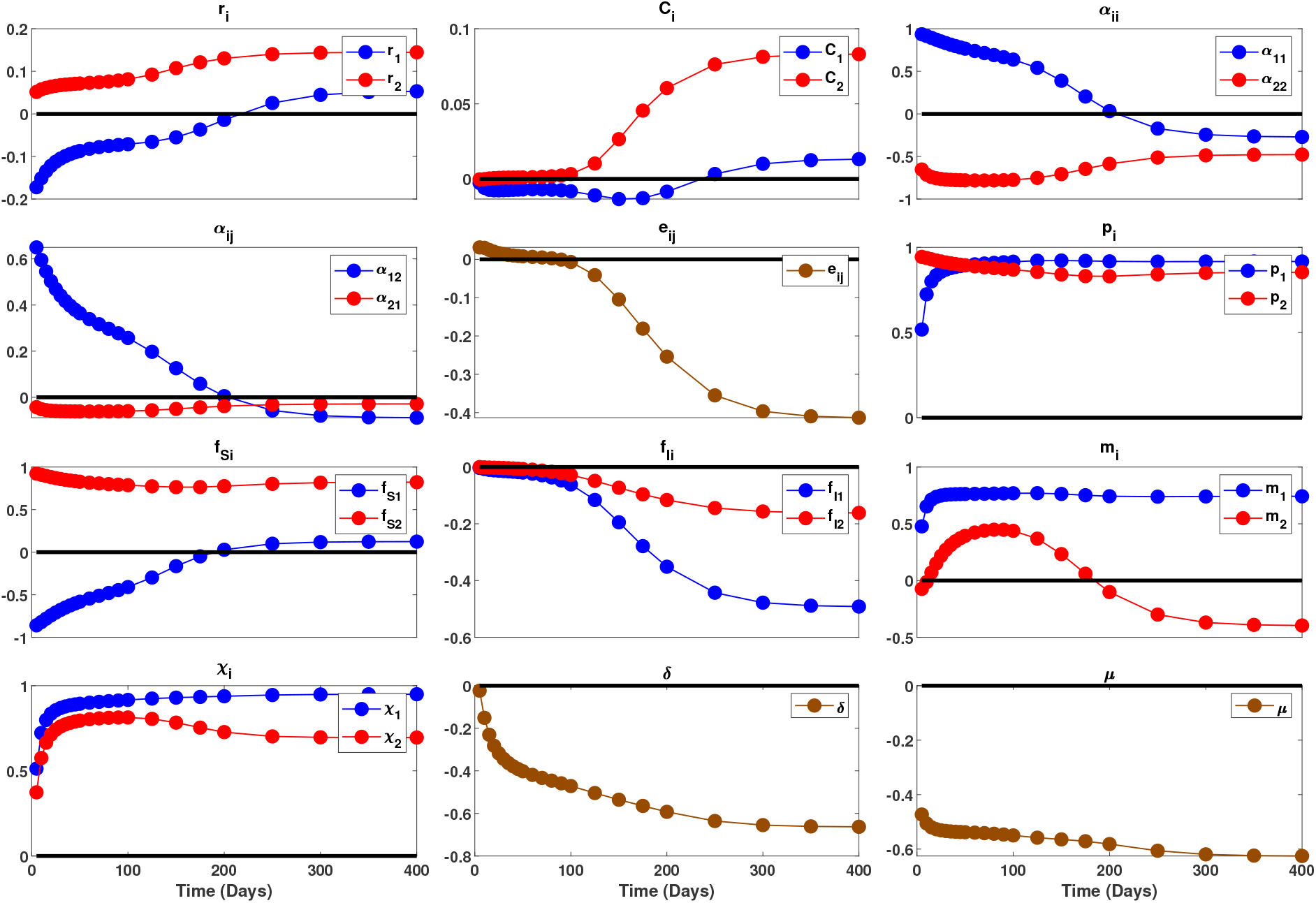
Global sensitivities of prevalence in host 2 (*I*_2_/*N*_2_) to all parameters in Case 1. In each panel, the blue curve corresponds to the PRCC values for a parameter for host 1, the red curve corresponds to a PRCC value for a parameter for host 2, and the black line denotes a value of zero.

## Susceptible host filtering rates 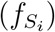

The sensitivity of infection prevalence in host *i* to the filtering rate in host *i* is positive for all time; this matches the intuition that increased transmission rates lead to increased prevalence. In comparison, the sensitivity of infection prevalence in host *i* to the filtering rate in host *j* (*j* ≠ *i*) changes from negative to positive. The sensitivity is initially negative because when spores are initially introduced, the rates of increase in infection prevalence in both hosts are limited by the low spore density. (The bottom panel of Figure B3 shows that spore density is much lower early in time than it is later in time for all simulations used to compute the PRCC sensitivity indices.) Higher filtering in host *j* removes spores that can potentially infect host *i*, which reduces the infection rate for host *i*. Later in time, the sensitivity becomes positive because spore density is higher and increased increased filtering by host *j* causes higher prevalence in host *j*, which causes greater release of spores by host *j*, which causes increased spore densities, which in turn results in more infections in host *i*.

**Figure B3:**
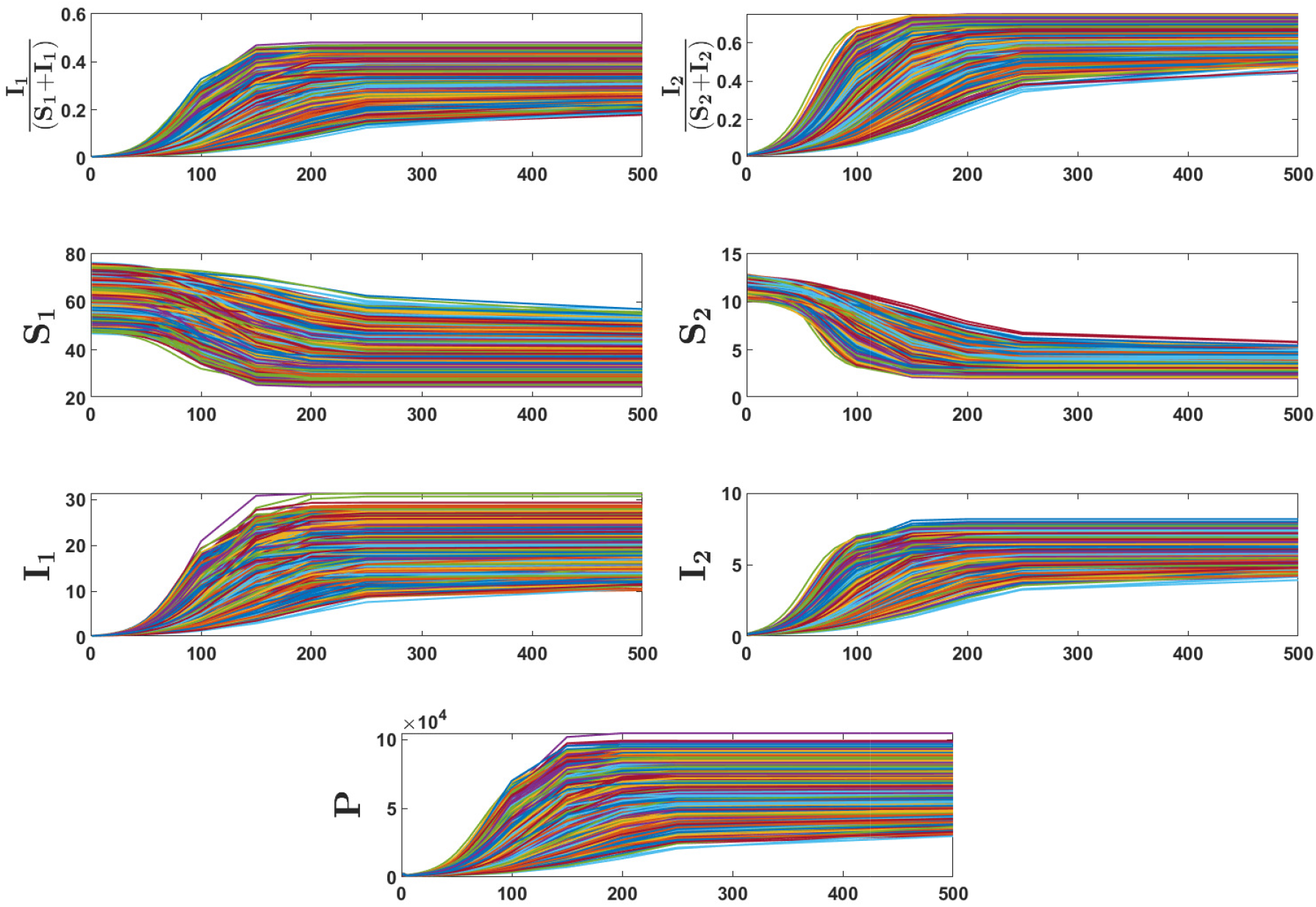
Plots of the time series used to compute the PRCC sensitivity indices for Case 1, where the pathogen is introduced into the disease-free two-host community at equilibrium. Colors correspond to individual simulations. Variation in the time series is due to variation in the parameter values. The bottom panel shows that the initial spore density is much lower than the densities at equilibrium for all parameter values.

## Host competition coefficients (*α_ij_*)

The sensitivities of the parameters defining competition on host 2 (*α*_21_, *α*_22_) are negative for both quantities of interest for all time points. This agrees with the intuition from the local sensitivities: increased competition reduces susceptible host density, which results in decreased infection rates and decreased infection prevalence. In contrast, the sensitivities of the parameters defining competition on host 1 (*α*_11_, *α*_12_) are positive for both quantities of interest for sufficiently small time. This positive relationship between competition on host 1 and disease prevalence is due to the initial condition of the system; see below for the mathematical justification. The biological reason is the following. When the spores are initially introduced, the rates of increase in infection prevalence in both hosts are limited by the low spore density. The second host produces spores faster than the first host because the second host has a higher transmission rate and a larger burst size 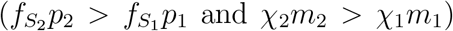. Consequently, the spore density will increase faster if more individuals of the second host become infected and fewer individuals of the first host become infected. Increasing competition on host 1 (i.e., increasing *α*_11_ or *α*_12_) causes a decrease in its equilibrium density and an increase in the equilibrium density of host 2. Thus, by increasing competition on host 1, spore density increases faster (via more infected individuals of host 2) and infection prevalence in both hosts increases faster. We note that the opposite reasoning is why competition on host 2 has a negative effect on host prevalence: increased competition on host 2 causes spore density to increase slower, which in turn causes infection prevalence to increase slower.

The mathematical explanation for the preceding paragraph is based on the local sensitivities of the derivatives *dY_i_*/*dt* evaluated near the disease-free equilibrium. Specifically, for sufficiently small *t*, the rate of change in infection prevalence of host *i* is approximately given by 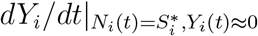 where 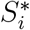 is the host density at disease free equilibrium. Note that 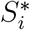 depends on *α_ij_* for *i, j* ∈ {1, 2}; see equations (9) and (10) in the main text. To leading order, the local sensitivities of the time derivatives to *α*_11_ are

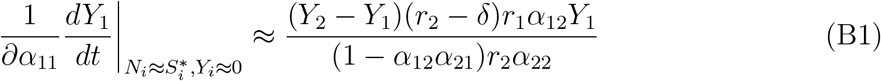

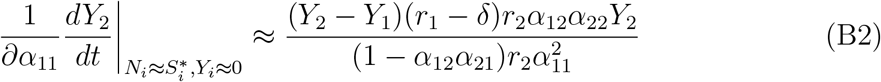

For the nominal parameter values, 1 − *α*_12_*α*_21_ > 0 and *r_i_* − *δ*> 0. This implies the signs of both local sensitivities are determined by *Y*_2_ − *Y*_1_. For sufficiently small *t*, *dY*_2_/*dt > dY*_1_/*dt* because to leading order 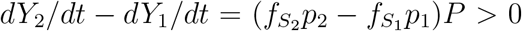, which means *Y*_2_(*t*) − *Y*_1_(*t*) > 0 for sufficiently small *t*. Consequently, the local sensitivities of the quantities of interest to *α*_11_ are positive. Nearly identical calculations show that the local sensitivities are positive for *α*_12_ and negative for *α*_21_ and *α*_22_.

## B.2 Case 2: Host 2 invades the endemic equilibrium of host 1

In Case 2, host 1 (the resident host) stably coexists with the pathogen at the endemic equilibrium and host 2 (the invading host) is introduced into the system. Time series for the global sensitivities for all parameters are given in Figures B4 and B5. For many of the most sensitive parameters 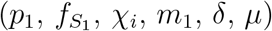, the signs of the global sensitivities at all points in time agree with the signs of the local sensitivities; see appendix A.2 for interpretations. However, differences in signs occur for some parameters of host 2 (*p*_2_, 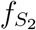, *m*_2_) over short times scales because total uptake of spores by host 2 is initially much greater than the total release of spores. Explanations for the differences are given below.

**Figure B4:**
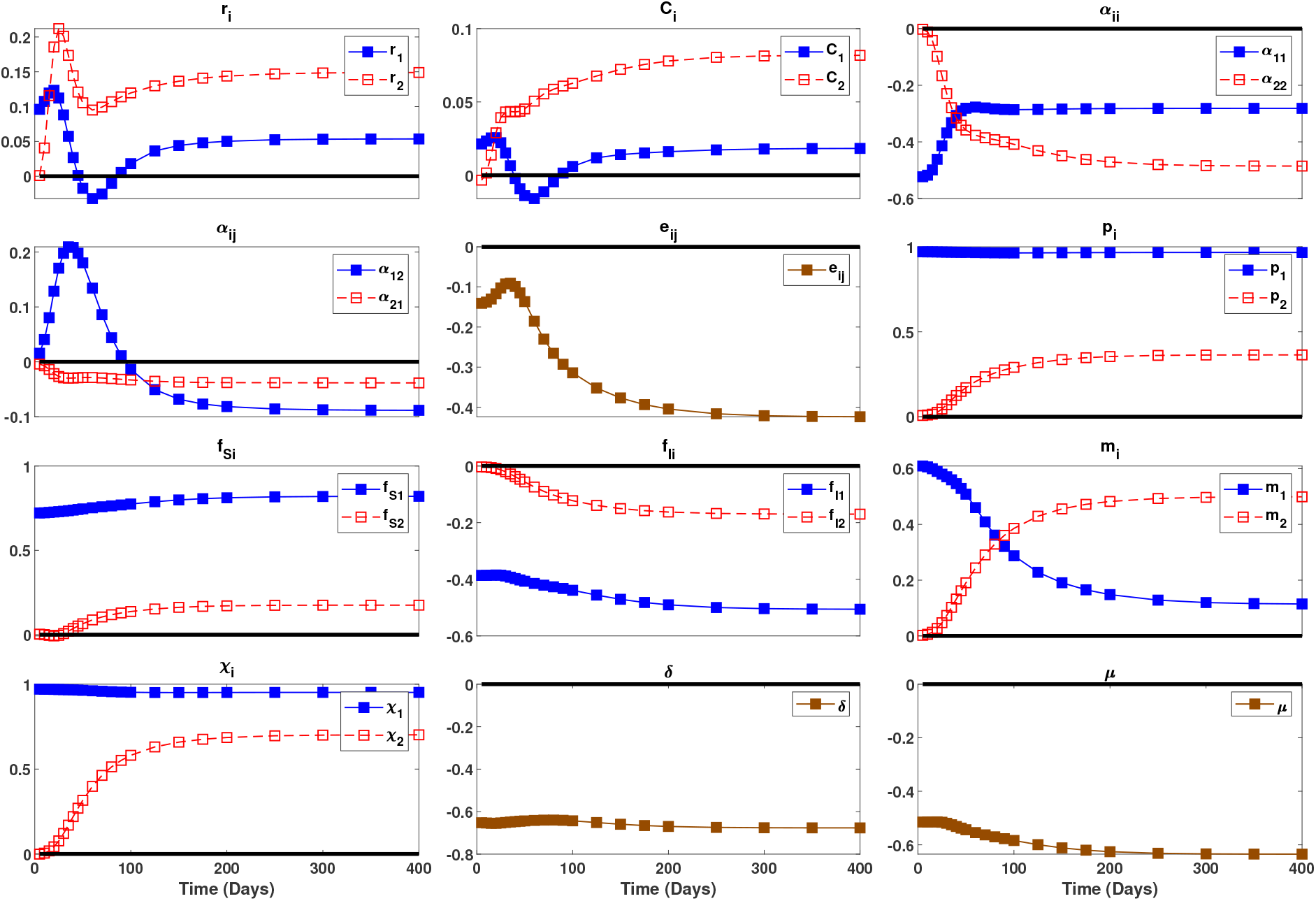
Global sensitivities of prevalence in host 1 (*I*_1_/*N*_1_) to all parameters in Case 2. In each panel, the solid blue curve with a filled symbols corresponds to the PRCC values for a parameter for host 1 (the resident), the dashed red curve with open symbols corresponds to a PRCC value for a parameter for host 2 (the invader), and the black line denotes a value of zero.

**Figure B5:**
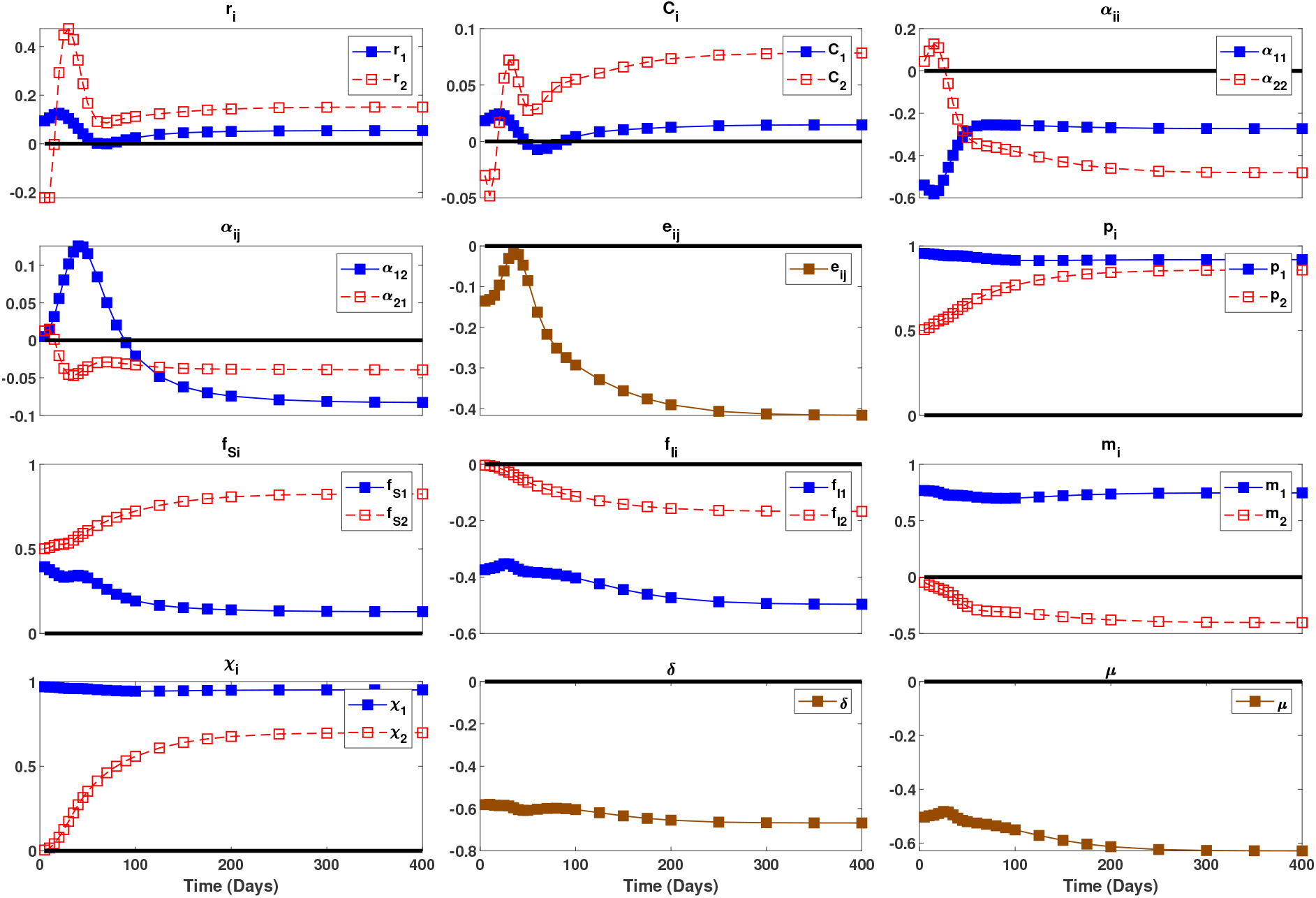
Global sensitivities of prevalence in host 1 (*I*_2_/*N*_2_) to all parameters in Case 2. In each panel, the solid blue curve with a filled symbols corresponds to the PRCC values for a parameter for host 1 (the resident), the dashed red curve with open symbols corresponds to a PRCC value for a parameter for host 2 (the invader), and the black line denotes a value of zero.

## Parameters affecting host infection rates (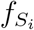, *p_i_*)

For most of the combinations of 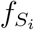 or *p_i_* and the quantities of interest, the global sensitivities are positive at all points in time; this matches the intuition from the local sensitivities. The two exceptions are that the local sensitivities of the resident host disease prevalence (*Y*_1_ = *I*_1_/*N*_1_) to the infection probability of host 2 (*p*_2_) and the uptake rate of susceptible individuals of host 2 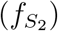 are initially negative and then become positive for sufficiently large time. For 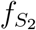, the reason is that increased filtering by host 2 decreases spore density and the decrease is not offset by spore release until later in time when infected individuals die. Consequently, infection rates for host 1 are initially lower when host 2 has higher filtering rates, but the infection rates ultimately become higher when many infected individuals of host 2 are releasing spores. For *p*_2_, the reason is that increased probability of infection by host 2 reduces competition on host 1. Specifically, when *p*_2_ is greater, a greater proportion of individuals in host 2 are infected and the total growth rate of host 2 (*dN*_2_/*dt*) is lower because infected individuals of host 2 have lower reproductive output than susceptible individuals of host 2 (*c*_2_ < 1). Fewer individuals in host 2 leads to less competition on host 1, which means the density of susceptible individuals decreases slower (Figure B6 shows the decrease in *S*_1_ immediately after the introduction of host 2 at time *t* = 1000). As a result, the proportion of susceptible individuals in host 1 decreases at a slower rate, which means the proportion of infected individuals of host 1 increases at a slower rate (the top left panel of Figure B6 shows the increase in *Y*_1_ = *I*_1_/*N*_1_ immediately after the introduction of host 2 at time *t* = 1000). Because of the slower increase in *Y*_1_, the sensitivity of prevalence in host 1 to *p*_2_ is negative.

**Figure B6:**
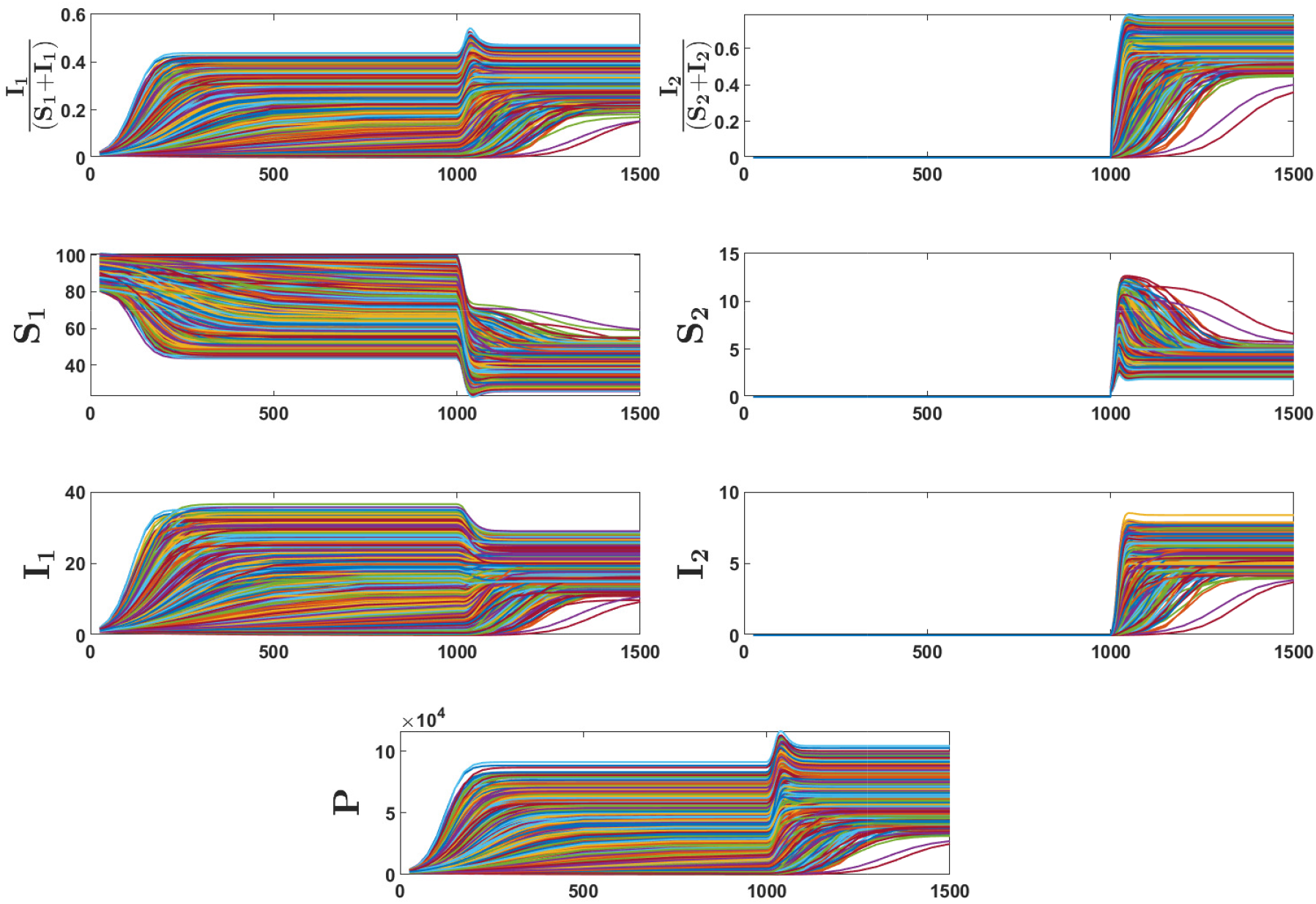
Plots of the time series used to compute the PRCC sensitivity indices for Case 2, where the host 2 is introduced into the endemic community made up of host 1 and the pathogen at equilibrium. Host 2 is introduced at *t* = 1000. Colors correspond to individual simulations. Variation in the time series is due to variation in the parameter values.

## Infected host mortality rates (*m_i_*)

Nearly all of the signs of the global sensitivities agree with the signs of the local sensitivities. The one exception is that the sensitivity of infection prevalence in host 1 to the mortality rate of host 2 (*m*_2_) transitions from negative to positive. The reason for the negative value is the following. Increased mortality of host 2 reduces its growth rate, leading to a slower increase in the density of host 2. This reduces the level of interspecific competition experienced by host 1, causing the density of susceptible individuals to decrease at a slower rate; see time series in Figure B6. Higher densities of susceptible individuals result in lower disease prevalence in host 1 (*Y*_1_ = *I*_1_/*N*_1_) with increased mortality of host 2.

## B.3 Case 3: Host 1 invades the endemic equilibrium of host 2

In Case 3, host 2 (the resident host) stably coexists with the pathogen at the endemic equilibrium and host 1 (the invading host) is introduced into the system. Time series for the global sensitivities for all parameters are given in Figures B7 and B8. For many of the most impactful parameters 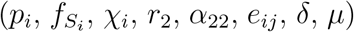, the signs of the global sensitivities at all points in time agree with the signs of the local sensitivities; see appendix A.2 for interpretations. However, differences in signs occur for the mortality rate of host 2 (*m*_2_) over short times scales because the infection rate of host 2 switches from being limited by spore density to being limited by the host density. In addition, differences in signs occur for some parameters of host 1 (*r*_1_, *α*_11_, *m*_1_) because increased mortality, increased reproduction, and reduced intraspecific competition initially cause the density of susceptible individuals to increase faster than the density of infected individuals. Explanations for the differences are given below.

**Figure B7:**
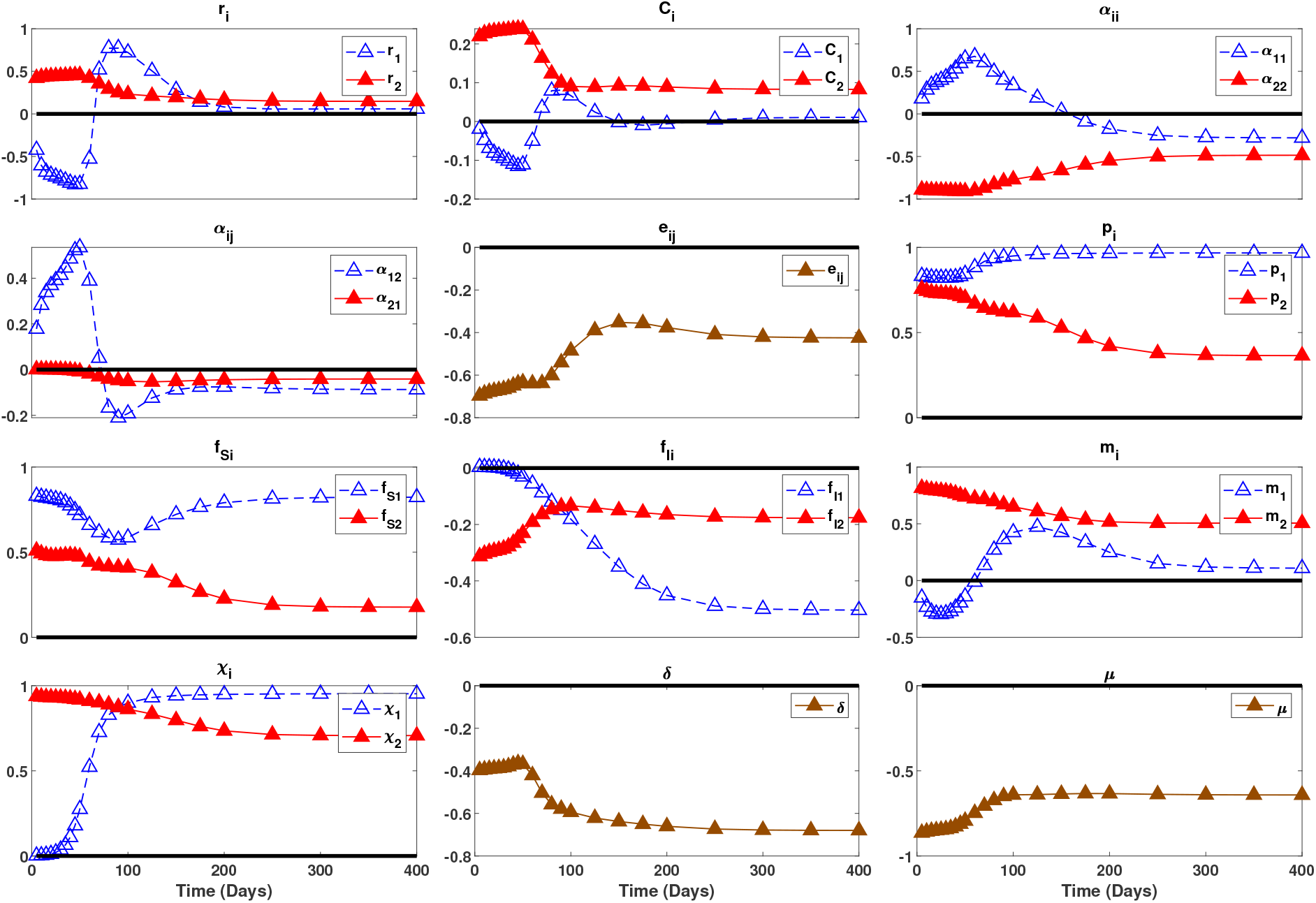
Global sensitivities of prevalence in host 1 (*I*_1_/*N*_1_) to all parameters in Case 3. In each panel, the dashed blue curve with a open symbols corresponds to the PRCC values for a parameter for host 1 (the invader), the solid red curve with filled symbols corresponds to a PRCC value for a parameter for host 2 (the resident), and the black line denotes a value of zero.

**Figure B8:**
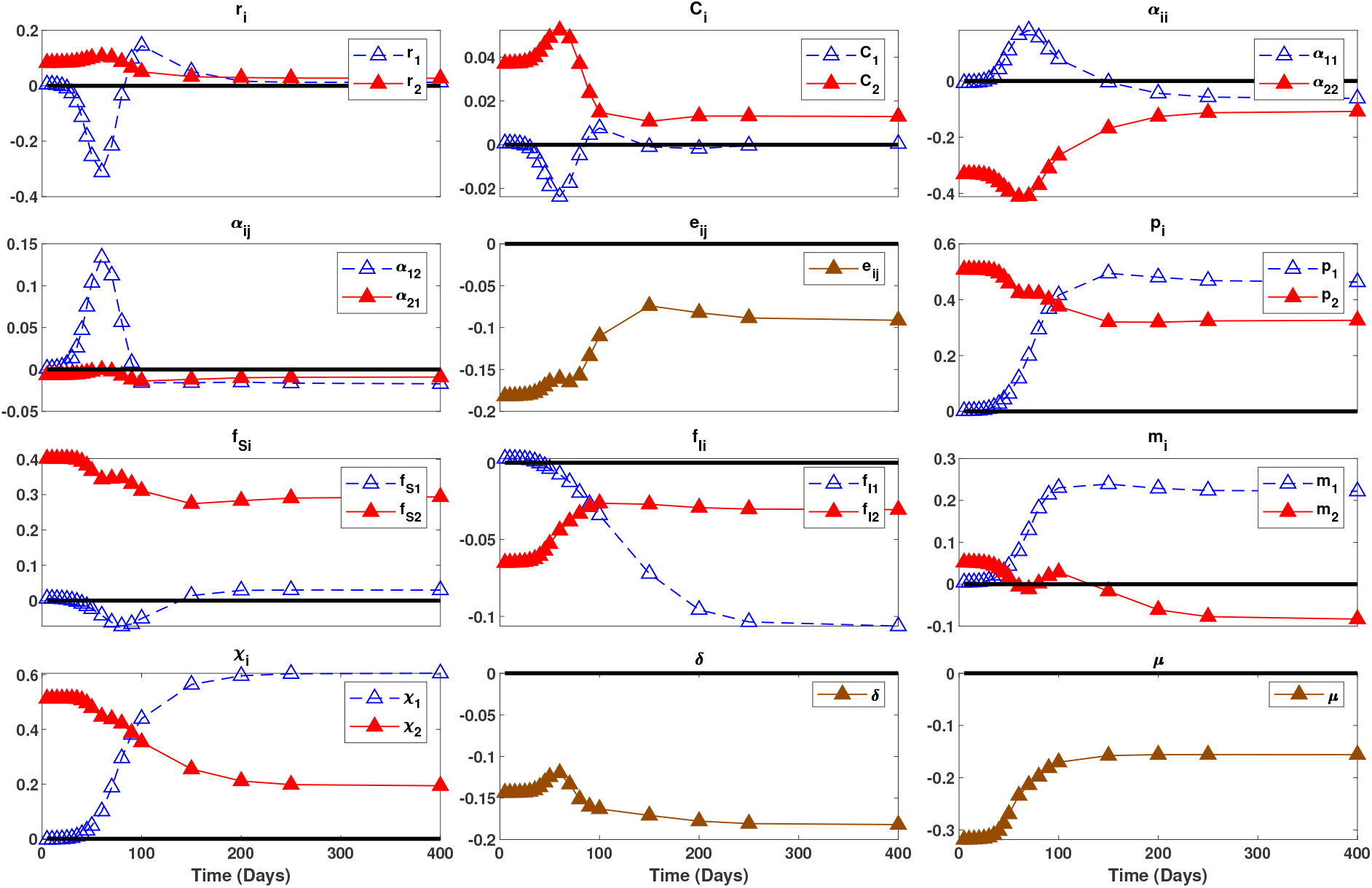
Global sensitivities of prevalence in host 1 (*I*_2_/*N*_2_) to all parameters in Case 3. In each panel, the dashed blue curve with a open symbols corresponds to the PRCC values for a parameter for host 1 (the invader), the solid red curve with filled symbols corresponds to a PRCC value for a parameter for host 2 (the resident), and the black line denotes a value of zero.

## Infected mortality rate for host 2 (*m*_2_)

Increased mortality of host 2 (*m*_2_) has a positive effect on infection prevalence in host 1 for all points in time; this matches the intuition from the local sensitivities. In contrast, the effects of increased mortality of host 2 (*m*_2_) on infection prevalence in host 2 change from negative to positive. This change in sign is driven by the transmission rate for host 2 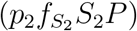. Early in time, the spore release rate of host 2 determines the spore density and spore density is the factor limiting the transmission rate for host 2. Because increased mortality of host 2 increases spore release rates and spore density, increased mortality of host 2 increases the transmission rate for host 2 and ultimately leads to increased infection prevalence in host 2. Later in time, the spore release rates of hosts 1 and 2 jointly determine the spore density and susceptible density of host 2 is the factor limiting the transmission rate for host 2. The negative effect of increased mortality on the density of host 2 (via reduced reproductive output by infected individuals) results in a negative effect of increased mortality on the transmission rate of host 2. As a result, increased mortality of host 2 causes decreased infection prevalence in host 2.

**Figure B9:**
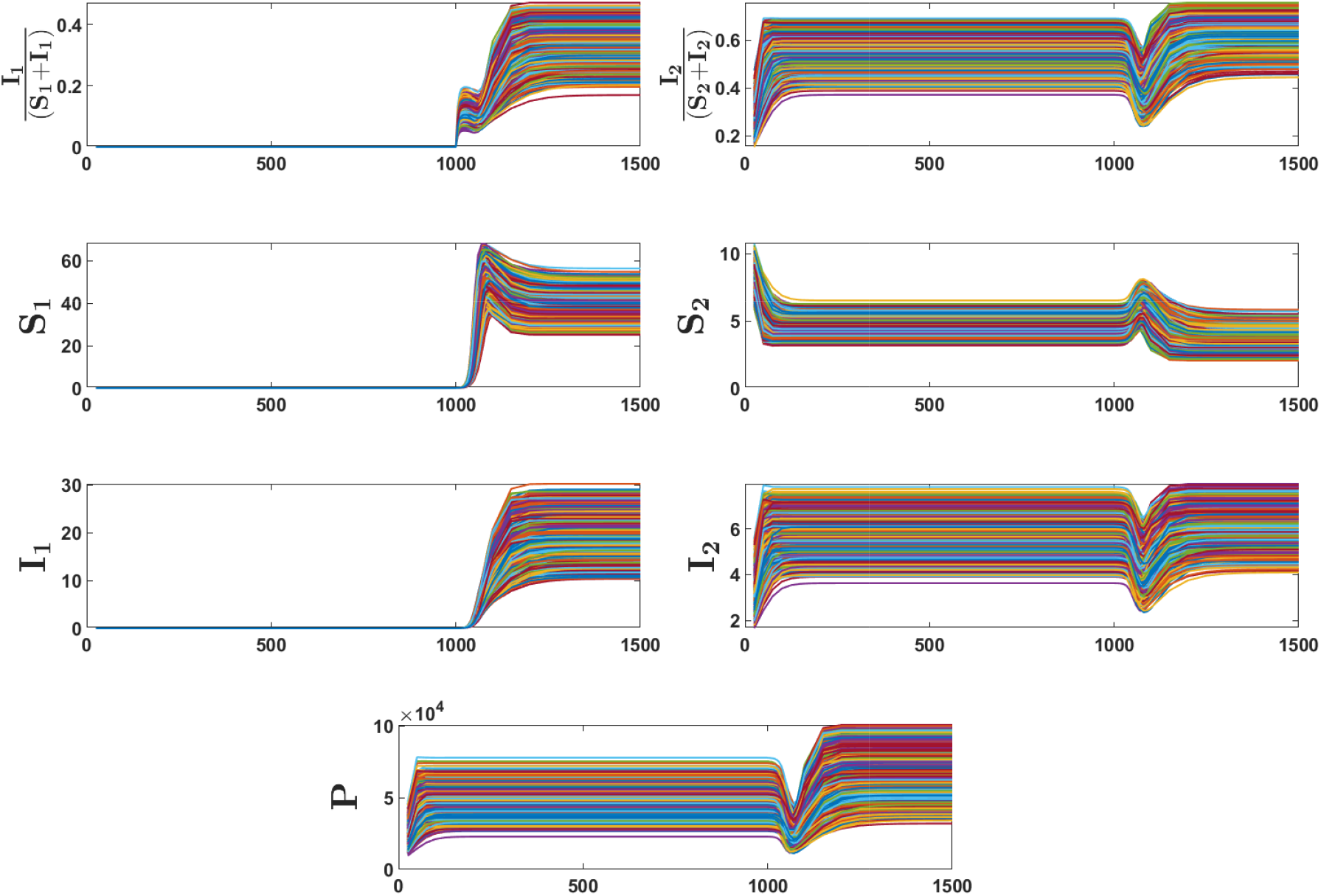
Plots of the time series used to compute the PRCC sensitivity indices for Case 3, where the host 1 is introduced into the endemic community made up of host 2 and the pathogen at equilibrium. Host 1 is introduced at *t* = 1000. Colors correspond to individual simulations. Variation in the time series is due to variation in the parameter values.

## Infected mortality rate (*m*_1_), reproduction rate (*r*_1_) and intraspecific competition coefficient (*α*_11_) for host 1

The sensitivity of infection prevalence in host 1 to the mortality rate of host 1 (*m*_1_) transitions from negative to positive; the transition is the same for the reproduction rate of host 1 (*r*_1_) and the opposite for the intraspecific competition coefficient (*α*_11_). In all three cases, the differences in sign early in time are driven by the relative rates of increase in susceptible and infected densities in host 1. Specifically, early in time when the density of host 1 is low, increased mortality of infected individuals, increased reproduction rates, and decreased intraspecific competition all result in relatively faster rates of increase in susceptible individuals and slower rates of increase in infected individuals. Consequently, early in time there are negative effects of increased mortality and reproduction and a positive effect of increased intraspecific competition on disease prevalence in host 1. Later in time when the density of host 1 is higher, increased mortality, increased reproduction rates, and decreased intraspecific competition lead to higher transmission rates because they yield higher spore densities and higher host densities. As a result, later in time there are positive effects of increased mortality and reproduction and a negative effect of increased intraspecific competition on disease prevalence in host 1.

The sensitivity of the infection prevalence in host 2 to the reproduction rate of host 1 (*r*_1_) transitions from positive to negative and back to positive; the sequence of signs is the opposite for the intraspecific competition coefficient (*α*_11_). The initial signs of the sensitivities are due to the effects of competition. Specifically, increased reproduction and decreased intraspecific competition increase the growth rate and density of host 1, which in turn decreases the density of susceptible individuals of host 2 due to interspecific competition. Because the decrease in susceptible density occurs without a change in infected density, infection prevalence in host 2 increases. The signs of the sensitivities change shortly after because of how host 1 affects spore density. In particular, increased reproduction and decreased intraspecific competition leads to higher densities of host 1, which causes greater decreases in in spore density and greater decreases the transmission rate of host 2. However, the signs of the sensitivities eventually switch back because infected individuals in host 1 reach sufficiently high densities that the release rate of spores by infected individuals of host 1 is greater than the total rate of uptake of spores by all host 1 individuals.

## C Detailed comparison of temporal dynamics across cases

The following provides detailed comparisons of the temporal dynamics of the global sensitivities across cases. Here, we focus on parameters whose PRCC values exceed 0.6 for at least one host at some point in time (i.e, max(|*PRCC*(*t*)|) ≥ 0.6). The Global sensitivities of each quantity of interest to the parameters are compared in Figures 6-8 in the main text.

## C.1 Detailed comparison of temporal dynamics in Case 1

Recall that in Case 1 the pathogen is introduced into the disease-free two-host community at equilibrium. The time series for the global sensitivities are plotted in Figure 6 of the main text.

## Probabilities of infection (*p_i_*)

Early in time, prevalence in each host species is more sensitive to its own probability of infection than the probability of infection of the other host (blue curve above red curve in top panel of column 2 for small *t* and red curve above blue curve in bottom panel of column 2 for small *t*). At the end of the simulation, both host species are more sensitive to the probability of infection for host 1 (*p*_1_; blue curves above red curves in both panels in column 2 for large *t*). The transient dynamics are non-monotonic for both parameters and both host species.

## Susceptible individual filtering rates 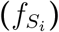

Early in time, prevalence in each host species is more sensitive to its own filtering rate than the filtering rate of the other host (blue curve is larger magnitude than the red curve in top panel of column 3 for small *t* and red curve is larger in magnitude than the blue curve in bottom panel of column 3 for small *t*). The same statement is true at the end of the simulation (blue curve above red curve in top panel of column 2 for large *t* and red curve above blue curve in bottom panel of column 2 for large *t*). The transient dynamics for the sensitivities of prevalence in host 1 are non-monotonic. Specifically, during the transient dynamics, prevalence in host 1 (*I*_1_/*N*_1_) becomes much more sensitive to the filtering rate of host 2 and much less sensitive to the filtering rate of host 1 (red curve much larger in magnitude than the blue curve in the top panel of column 2 for *t* ≈ 75). In comparison, the transient dynamics for the sensitivities of prevalence in host 2 (*I*_2_/*N*_2_) are nearly non-monotonic.

## Infected host mortality rates (*m_i_*)

Early in time, prevalence in each host species is more sensitive to the mortality rate of the other host than its own filtering rate (red curve is larger magnitude than the blue curve in top panel of column 3 for small *t* and blue curve is larger in magnitude than the red curve in bottom panel of column 4 for small *t*). The same statement is true at the end of the simulation (red curve is larger magnitude than the blue curve in top panel of column 3 for large *t* and blue curve is larger in magnitude than the red curve in bottom panel of column 4 for large *t*). The transient dynamics for the sensitivities of both parameters are non-monotonic. Interestingly, the sensitivity of prevalence in host 2 to its own mortality rate changes from negative, to positive, back to negative (see bottom panel of column 4). The negative signs for small and large *t* are likely due to host 2’s small population size. We hypothesize that the positive sign for intermediate time is due to a positive feedback: increased mortality leads to increased release of spores, which then leads to greater infection rates in host 2. This positive feedback is sufficiently strong at intermediate time because the population is exponentially growing, which leads to an abundance of susceptible individuals that can become infected.

## Burst sizes (*χ_i_*)

At all points in time prevalence in both host species is more sensitive to the spore burst size of host 1 than host 2 (blue curves above red curves in both panels of column 5).

## Intraspecific competition coefficients (*α_ii_*)

Early in time, prevalence in each host species is more sensitive to the intraspecific competition coefficient for host 1 than host 2 (blue curves larger in magnitude than red curves in both panels of column 1 for small *t*). In addition, the sensitivity is positive for the intraspecific competition coefficient for host 1 and negative for the intraspecific competition coefficient for host 2; see equations B1 and B2 and surrounding text for an explanation for the difference in sign. As explained in the main text, we hypothesize that this is due to the difference in disease competence for the two host species. At the end of the simulation, both host species are more sensitive to the intraspecific competition coefficient for host 2 (*α*_22_; red curves larger in magnitude than red curves in both panels of column 2 for large *t*). During the transient dynamics, the sensitivities to the competition coefficient for host 2 monotonically decrease from large positive values to small negative values. In contrast, the sensitivities to the competition coefficient for host 2 are u-shaped. Thus, prevalence in both host species becomes less sensitive to intraspecific competition in host 1 but transiently becomes more sensitive to intraspecific competition in host 2.

## C.2 Detailed comparison of temporal dynamics in Cases 2 and 3

Recall that in Cases 2 and 3, one host is introduced into the community where the other host and pathogen coexist at their endemic equilibrium. The time series for the global sensitivities are plotted in Figures 8 and 8 of the main text.

## Probabilities of infection (*p_i_*)

Early in time, prevalence in both host species is more sensitive to the resident’s probability of infection than the invader’s probability of infection (solid curves with a given shape above dashed curves of the opposite color with the same shape in column 1 of figure 7). At the end of the simulation, prevalence in both host species is more sensitive to the infection probability of host 1 than that of host 2. The changes in most sensitivities are monotonic. The only exception to the above is in Case 3 where host 1 (the invader) is slightly more sensitive to its probability of infection than that of host 2 (the resident) early in time (dashed blue triangles above solid red triangles in top panel of column 1 in Figure 8) and the change in the sensitivity over time is nearly monotonic (see *t* = 50).

## Susceptible individual filtering rates 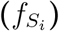

For all points in time, prevalence in each host species is more sensitive to its filtering rate than that of the other host species (blue curves above red curves in top panel of column 2 of figure 7 and red curves above blue curves in bottom panel of column 2 of figure 7). The transient dynamics are monotonic or nearly monotonic for most sensitivities; the exception is the non-monotonic changes in the sensitivities of prevalence in both host species to the filtering rate of host 1 (the invader) in Case 3.

## Infected host mortality rates (*m_i_*)

Early in time, prevalence in both host species is more sensitive to the mortality rate of the resident host than the invading host (solid curves larger in magnitude than dashed curves in both panels of column 3 of Figure 7 for small *t*). At the end of the simulation, prevalence in each host species is more sensitive to the mortality rate of the other host than its own mortality rate (red curves above blue curves in top panel of column 3 of Figure 7 for small *t* and blue curves above red curves in bottom panel of column 3 of Figure 7 for small *t*). The transient dynamics are nearly monotonic for Case 2. For Case 3, the sensitivity of prevalence in each host to its own mortality rate is nonlinear (dashed blue triangles in top panel of column 3 of Figure 7 and solid red triangles in bottom panel of column 3 of Figure 7). The authors are unaware of clear explanation for the non-monotonic changes in the sensitivities.

## Spore burst size rates (*χ_i_*)

Early in time, prevalence in both host species is more sensitive to the resident’s burst size than the invader’s burst size (solid curves with a given shape above dashed curves of the opposite color with the same shape in column 4 of Figure 7). At the end of the simulation, prevalence in both host species is more sensitive to the burst size of host 1 than host 2 (blue curves above red curves in both panels of column 4 of Figure 7). The sensitivities for both host species change monotonically.

## Maximum exponential growth rates (*r_i_*)

The sensitivities of prevalence in both host species to the exponential growth rates of the species are small in magnitude for both Cases 2 and 3. The only exception is the sensitivity to the exponential growth rate of host 1 (the invader) in Case 3 (dashed curves in both panels of column 1 of Figure 8); for both host species, the sensitivity decreases to a large negative value, increases to a large positive value and then ultimately converges to small positive value. We hypothesize that this non-monotonic change is due to the non-monotonic dynamics of the susceptible class (*S*_1_ time series in Figure B9). In particular, early in time (*t* ≈ 1000 in Figure B9), prevalence in host 1 increases exponentially and the host growth rate (*r*_1_) has a negative sensitivity value because faster growth of susceptible individuals yields a lower growth in prevalence. Later in time (*t* ≈ 1075 in Figure B9), the susceptible class for host 1 (*S*_1_) overshoots its equilibrium density and begins to decrease. Here, the host growth rate (*r*_1_) has a positive sensitivity value because faster growth of susceptible individuals would increase the density of susceptible individuals, which results in a greater density of infected individuals.

## Intraspecific competition coefficients (*α_ii_*)

Early in time, prevalence in each host species is more sensitive to the intraspecific competition coefficient for the resident host than the invading host (solid curves with a given shape are larger in magnitude than dashed curves with the same shape and of the opposite color in both panels of column 2 of Figure 8 for small *t*). At the end of the simulation, both host species are more sensitive to the intraspecific competition coefficient for host 2 (*α*_22_; red curves larger in magnitude than red curves in both panels of column 2 of Figure 8 for large *t*). The transient dynamics of the sensitivities are non-monotonic in both cases, but larger variation is observed in Case 3. In addition, the sensitivities to the invader’s parameter transition from negative to positive back to negative whereas the sensitivities to the resident’s parameter are always negative. The authors are unaware of a clear explanation for why these differences occur between the cases and between the resident’s and invader’s traits.

## Interspecific competition coefficients (*α_ij_*)

For both cases, the sensitivities to host 1’s interspecific competition coefficient (*α*_21_) are small in magnitude (blue curves small in magnitude in both panels of column 4 of Figure 8). In contrast, the sensitivities to host 2’s interspecific competition coefficient (*α*_12_) are large in magnitude for finite periods of time (red curves in both panels of column 4 of Figure 8). In addition, the signs of the sensitivities to *α*_12_ change from positive to negative. The reason for the sign change is the same as that for intraspecific competition coefficients (after accounting for the fact that the interspecific competition coefficient for host *i*, *α_ji_*, affects the growth rate of host *j* where *j* ≠ *i*).

## References

Aggarwal, M., N. Cogan, and R. Bertram. 2019. Where to look and how to look: Combining global sensitivity analysis with fast/slow analysis to study multi-timescale oscillations. Mathematical biosciences 314:1–12.

Aggarwal, M., N. Cogan, and O. L. Lewis. 2021. Physiological insights into electrodiffusive maintenance of gastric mucus through sensitivity analysis and simulations. Journal of Mathematical Biology 83:1–31.

Becker, C. G., D. Rodriguez, L. F. Toledo, A. V. Longo, C. Lambertini, D. T. Corrêa, D. S. Leite, C. F. Haddad, and K. R. Zamudio. 2014. Partitioning the net effect of host diversity on an emerging amphibian pathogen. Proceedings of the Royal Society B: Biological Sciences 281:20141796.

Blower, S. M., and H. Dowlatabadi. 1994. Sensitivity and uncertainty analysis of complex models of disease transmission: an hiv model, as an example. International Statistical Review/Revue Internationale de Statistique pages 229–243.

Cortez, M. H. 2021. Using sensitivity analysis to identify factors promoting higher versus lower infection prevalence in multi-host communities. Journal of Theoretical Biology 526:110766.

Cortez, M. H., and M. A. Duffy. 2021. The context-dependent effects of host competence, competition, and pathogen transmission mode on disease prevalence. The American Naturalist 198:000–000.

Dizney, L. J., and L. A. Ruedas. 2009. Increased host species diversity and decreased prevalence of Sin Nombre Virus. Emerging Infectious Diseases 15:1012–1018.

Dobson, A. 2004. Population dynamics of pathogens with multiple host species. American Naturalist 164:S64–S78.

Evans, H. F., and P. F. Entwistle. 1987. Viral diseases. In J. R. Fuxa and T. Tanada, eds., Epizootiology of Insect Diseases. Wiley, New York.

Faust, C. L., A. P. Dobson, N. Gottdenker, L. S. Bloomfield, H. I. McCallum, T. R. Gillespie, M. Diuk-Wasser, and R. K. Plowright. 2017. Null expectations for disease dynamics in shrinking habitat: dilution or amplification? Philosophical Transactions of the Royal Society B: Biological Sciences 372:20160173.

Halliday, F. W., J. R. Rohr, and A.-L. Laine. 2020. Biodiversity loss underlies the dilution effect of biodiversity. Ecology letters 23:1611–1622.

Hamby, D. 1995. A comparison of sensitivity analysis techniques. Health physics 68:195–204.

Hanthanan Arachchilage, K., and M. Hussaini. 2021. Ranking non-pharmaceutical interventions against covid-19 global pandemic using global sensitivity analysis-effect on number of deaths. Chaos, Solitons & Fractals 152:111458.

Hopkins, S. R., A. E. Fleming-Davies, L. K. Belden, and J. M. Wojdak. 2020. Systematic review of modelling assumptions and empirical evidence: Does parasite transmission increase nonlinearly with host density? Methods in Ecology and Evolution 11:476–486.

Hydeman, M. E., A. V. Longo, G. Velo-Antón, D. Rodriguez, K. R. Zamudio, and R. C. Bell. 2017. Prevalence and genetic diversity of *batrachochytrium dendrobatidis* in Central African island and continental amphibian communities. Ecology and Evolution 7:7729–7738.

Jansen, M. 1999. Analysis of variance designs for model output. Computer Physics Communications.

Jarrett, A., N. Cogan, and M. Hussaini. 2017a. Combining two methods of global sensitivity analysis to investigate mrsa nasal carriage model. Bulletin of Mathematical Biology 79:2258–2272.

Jarrett, A. M., N. Cogan, and M. Hussaini. 2017b. Combining two methods of global sensitivity analysis to investigate mrsa nasal carriage model. Bulletin of mathematical biology 79:2258–2272.

Jarrett, A. M., Y. Liu, N. Cogan, and M. Y. Hussaini. 2015. Global sensitivity analysis used to interpret biological experimental results. Journal of Mathematical Biology 71:151–170.

Johnson, P. T., R. B. Hartson, D. J. Larson, and D. R. Sutherland. 2008. Diversity and disease: community structure drives parasite transmission and host fitness. Ecology Letters 11:1017–1026.

Joseph, M. B., J. R. Mihaljevic, S. A. Orlofske, and S. H. Paull. 2013. Does life history mediate changing disease risk when communities disassemble? Ecology Letters 16:1405–1412.

Kucherenko, S., S. Tarantola, and P. Annoni. 2012. Estimation of global sensitivity indices for models with dependent variables. Computer Physics Communications 183:937–946.

Levine, R. S., D. L. Hedeen, M. W. Hedeen, G. L. Hamer, D. G. Mead, and U. D. Kitron. 2017. Avian species diversity and transmission of west nile virus in Atlanta, Georgia. Parasites & Vectors 10:62.

LoGiudice, K., S. T. Duerr, M. J. Newhouse, K. A. Schmidt, M. E. Killilea, and R. S. Ostfeld. 2008. Impact of host community composition on lyme disease risk. Ecology 89:2841–2849.

Luis, A. D., A. J. Kuenzi, and J. N. Mills. 2018. Species diversity concurrently dilutes and amplifies transmission in a zoonotic host–pathogen system through competing mechanisms. Proceedings of the National Academy of Sciences 115:7979–7984.

Mara, T., and S. Tarantola. 2012. Variance-based sensitivity indices for models with dependent inputs. Reliability Engineering and System Safety 107:115–121.

Marino, S., I. Hogue, C. Ray, and D. Kirschner. 2008. A methodology for performing global uncertainty and sensitivity analysis in systems biology. Journal of Theoretical Biology 254:178–196.

Mihaljevic, J. R., M. B. Joseph, S. A. Orlofske, and S. H. Paull. 2014. The scaling of host density with richness affects the direction, shape, and detectability of diversity-disease relationships. PLOS One 9:e97812.

O’Regan, S. M., J. E. Vinson, and A. W. Park. 2015. Interspecific contact and competition may affect the strength and direction of disease-diversity relationships for directly transmitted microparasites. The American Naturalist 186:480–494.

Orlofske, S. A., R. C. Jadin, D. L. Preston, and P. T. Johnson. 2012. Parasite transmission in complex communities: predators and alternative hosts alter pathogenic infections in amphibians. Ecology 93:1247–1253.

Randolph, S. E., and A. D. M. Dobson. 2012. Pangloss revisited: a critique of the dilution effect and the biodiversity-buffers-disease paradigm. Parasitology 139:847–863.

Renardy, M., C. Hult, S. Evans, J. J. Linderman, and D. E. Kirschner. 2019. Global sensitivity analysis of biological multiscale models. Current opinion in biomedical engineering 11:109–116.

Roberts, M., and J. Heesterbeek. 2018. Quantifying the dilution effect for models in ecological epidemiology. Journal of The Royal Society Interface 15:20170791.

Roche, B., A. P. Dobson, J. F. Guegan, and P. Rohani. 2012. Linking community and disease ecology: the impact of biodiversity on pathogen transmission. Philosophical Transactions of the Royal Society B 367:2807–2813.

Rohr, J. R., D. J. Civitello, F. W. Halliday, P. J. Hudson, K. D. Lafferty, C. L. Wood, and E. A. Mordecai. 2020. Towards common ground in the biodiversity–disease debate. Nature Ecology & Evolution 4:24–33.

Rudolf, V. H. W., and J. Antonovics. 2005. Species coexistence and pathogens with frequency-dependent transmission. American Naturalist 166:112–118.

Salelli, A., S. Tarantola, F. Campolongo, and M. Ratto. 2004. Sensitivity Analysis in Practice, a guide to assessing scientific models. John Wiley & Sons Ltd.

Saltelli, A. 2002. Sensitivity analysis for importance assessment. Risk analysis 22:579–590.

Saltelli, A., K. Aleksankina, W. Becker, P. Fennell, F. Ferretti, N. Holst, S. Li, and Q. Wu. 2019. Why so many published sensitivity analyses are false: A systematic review of sensitivity analysis practices. Environmental Modeling and Software 114:29–39.

Saltelli, A., M. Ratto, S. Tarantola, and F. Campolongo. 2005. Sensitivity analysis for chemical models. Chemical reviews 105:2811–2828.

Searle, C. L., L. M. Biga, J. W. Spatafora, and A. R. Blaustein. 2011. A dilution effect in the emerging amphibian pathogen *Batrachochytrium dendrobatidis*. Proceedings of the National Academy of Sciences 108:16322–16326.

Searle, C. L., M. H. Cortez, K. K. Hunsberger, D. C. Grippi, I. A. Oleksy, C. L. Shaw, S. B. de la Serna, C. L. Lash, K. L. Dhir, and M. A. Duffy. 2016. Population density, not host competence, drives patterns of disease in an invaded community. American Naturalist 188:554–566.

Sobol, I. 2001. Global sensitivity indices for nonlinear mathematical models and their monte carlo estimates. Mathematics and Computers in Simulations 55:271–280.

Strauss, A. T., D. J. Civitello, C. E. Cáceres, and S. R. Hall. 2015. Success, failure and ambiguity of the dilution effect among competitors. Ecology Letters 18:916–926.

Telfer, S., K. Bown, R. Sekules, M. Begon, T. Hayden, and R. Birtles. 2005. Disruption of a host-parasite system following the introduction of an exotic host species. Parasitology 130:661.

Venesky, M. D., X. Liu, E. L. Sauer, and J. R. Rohr. 2014. Linking manipulative experiments to field data to test the dilution effect. Journal of Animal Ecology 83:557–565.

Wentworth, M., R. Smith, and H. Banks. 2016. Parameter selection and verification techniques based on global sensitivity analysis illustrated for an hiv model. SIAM/ASA Journal of Uncertainty Quantification 4:266–297.

Wu, Y.-T. 1994. Computational methods for efficient structural reliability and reliability sensitivity analysis. AIAA journal 32:1717–1723.

Zimmermann, M. R., K. E. Luth, and G. W. Esch. 2017. Snail species diversity impacts the infection patterns of echinostoma spp.: examples from field collected data. Acta Parasitologica 62:493–501.

